# Lack of centrosomes limits the epithelial-to-mesenchymal transition in human mammary cells

**DOI:** 10.64898/2026.09.18.752689

**Authors:** Maria Lucia Pigazzini, Johanna Melisa Tocci, Mathias Boulanger, Ferris Jung, Vladimir Benes, Ingrid Hoffmann, Niccolò Banterle

## Abstract

As the main microtubule network organisers of animal cells, centrosomes orchestrate diverse cellular processes. Centrosomes have been linked to cancer when numerically or structurally aberrant, yet their precise roles at different cancer stages remains to be defined. Here, we study centrosome function in the epithelial-to-mesenchymal transition (EMT), a key driver of cancer initiation, progression and metastasis. We first depleted centrosomes in a breast cancer mesenchymal-like model, resulting in insignificant cellular or nuclear rearrangements. We then investigated non-carcinogenic breast cells undergoing the EMT via TGFβ signalling, and removed the centrosomes before triggering this transition. Stimulated epithelial cells acquired a mesenchymal phenotype and rearranged their cytoskeleton. By contrast, stimulated cells lacking centrosomes retained epithelial morphology, showed little cytoskeletal remodelling, yet still altered nuclear shape and upregulated key EMT markers. Transcriptome analysis revealed reduced expression of important genes involved in extracellular matrix organisation and a downregulation of cell junction components when lacking centrosomes; further analysis revealed paxillin showing reduced colocalisation with actin focal adhesions. Centrosomes thus prove essential for cytoskeletal remodelling in the EMT of breast cancer cells.

## Introduction

In animal cells, centrosomes fulfil the function of main microtubule organising centres (MTOCs). Centrosomes are megadalton membrane-less cytoplasmic organelles comprising a pair of microtubule-based barrel-like structures called centrioles. Centrioles are decorated with dozens of interacting proteins, and surrounded by a crowded and dynamic protein matrix, the pericentriolar material (PCM) (Banterle & Gönczy, 2017). Centrosomes nucleate microtubules (MTs), which are responsible for critical cellular processes including: organising the cytoskeletal architecture, establishing polarity, supporting movement and serving as signalling hubs (Bornens, 2012). Importantly, via their role in assisting bipolar spindle formation, centrosomes ensure that the genomic content of the cell is inherited equally and faithfully by each daughter after cell division (Hoffmann, 2021). Due to the multifaceted and fundamental nature of their functions, centrosomes have been implicated in a variety of diseases, from ciliopathies to neurodevelopmental and neurodegenerative disorders, and centrosome abnormalities are emerging as a hallmark of cancer (Goundiam & Basto, 2021). Structural and numerical aberrations - from supernumerary centrioles to disrupted structure and length, and atypical PCM composition - have been described in a range of hematological and solid cancers (Lingle *et al*, 1998; Marteil *et al*, 2018; Köhrer *et al*, 2023). As an oncogenic mechanism, amplified centrosomes lead to chromosomal missegregation and chromosomal instability, increasing the risk of further mutations and adding survival advantages to cancer cells (Ganem *et al*, 2009; Godinho & Basto, 2025). Furthermore, the master regulator of centrosome biogenesis polo-like kinase 4 (PLK4) is misregulated in a variety of cancers, correlating with disease severity and poor clinical outcomes (Mu *et al*, 2022); and PLK4 overexpression has been shown experimentally to drive spontaneous tumorigenesis in mice and flies (Basto *et al*, 2008; Gönczy, 2015; Levine *et al*, 2017). It has thus become increasingly clear that centrosome abnormalities profoundly impact cancer biology. However, alongside the well described centriolar overexpression phenotypes, seemingly paradoxically, centrosome loss has also been found in prostate, pancreatic and ovarian cancer, occasionally even more frequently than centrosome excess within the same tissue, highlighting that both aberrancies can also coexist (Morretton *et al*, 2019; Wang *et al*, 2020; Morretton *et al*, 2022a). It is however still unclear how widespread centrosome loss actually is and what are its pathophysiological effects and consequences in general and at different stages of cancer initiation and progression (Kalbfuss & Gönczy, 2023). Given the fundamental role of PLK4 in centriole duplication as well as cell migration and division, this kinase emerged as a prime target for cancer therapeutics and several pharmacological agents are currently being clinically trialled (Holland & Cleveland, 2014; Parsyan *et al*, 2025).

The epithelial-to-mesenchymal transition (EMT) is a physiological programme that drives mammary gland development, tissue remodelling and wound healing, becoming hijacked during breast cancer initiation, progression and metastasis (Hanahan, 2022; Huang et al, 2022). As the most common malignancy in women worldwide, breast cancer remains a leading cause of cancer-related mortality, largely due to metastatic disease (Dillekås *et al*, 2019; Xiong *et al*, 2025; Bhangdia *et al*, 2026). The EMT is now recognized as a continuum of epithelial, hybrid and mesenchymal cellular states rather than a binary switch, with cells progressively losing epithelial polarity and adhesion, while acquiring migratory and invasive properties (Pastushenko *et al*, 2018; Williams *et al*, 2019; Yang *et al*, 2020);(Lamouille *et al*, 2014; Nieto *et al*, 2016). The morphology change, from cuboidal to spindle-like, alongside a signature shift in the cellular transcription and translation profile, as for example increase in SNAIL family transcription factors and decrease in E-cadherin adhesion, are the gold standard for identifying the EMT (Yilmaz & Christofori, 2009). Degradation of the basal membrane and an extensive remodelling of the underlying extracellular matrix (ECM) further promote an invasive phenotype (Sleeboom *et al*, 2024). One of the principal drivers of the EMT is cytokine transforming growth factor beta (TGFβ) signalling (Massagué & Sheppard, 2023). TGFβ activates EMT transcriptional programmes including promoting extensive remodelling of the cytoskeletal components (Datta et al, 2021; Lim et al, 2026): actin filaments and their actin-binding protein (Melchionna *et al*, 2021), microtubules and their networks (Pimm & Henty-Ridilla, 2021; Nurmagambetova *et al*, 2023) and intermediate filament (Romet-Lemonne *et al*, 2025). Consistent with these profound changes in cell architecture, previous studies have shown that centrosomes undergo dynamic repositioning during the EMT, suggesting an active role in establishing the mesenchymal state (Burute *et al*, 2017). However, whether centrosomes merely respond to the transition or actively coordinate the structural and molecular programmes underlying TGFβ-driven EMT remains unclear.

In this study, we explored the contribution of centrosomes, and the effects of their removal, to the EMT and the initiation of cancer, with special emphasis on how they impact the cytoskeleton. We employed a variety of imaging techniques coupled with super-resolution light-microscopy and subsequent image analysis in mesenchymal-like as well as non-carcinogenic human mammary cells. We further performed transcriptomic analysis to confirm how lack of centrosome deregulates cellular pathways active in cells undergoing the EMT and cancer initiation, and to identify key players of the actin network related to this transition. Our work in human breast cells uncovers a pathway for the lack of centrosomes to limit the EMT, possibly imposing an alternate, intermediate, state of this reprogramming; and highlights a potentially necessary role of centrosomes in the initiation of pathology.

## Results

### Centrosome removal does not affect mesenchymal-like BC cells

We first wondered whether centrosomes are necessary for the maintenance of the EMT programme and depleted them in the triple negative breast cancer (TNBC) cell line MDA-MB-231 with the potent and selective small molecule PLK4 inhibitor Centrinone B (CenB) (Wong *et al*, 2015). CenB acts by reversibly binding to PLK4 and preventing formation of the centriolar cartwheel, thus effectively blocking the centriolar duplication cycle (Habedanck *et al*, 2005; Holland *et al*, 2012). MDA-MB-231 is a well characterised BC cell line, with clear mesenchymal features, including spindle-like morphology, lack of contact and colony formation, and with heightened metastatic potential (Prat *et al*, 2010; Harrell *et al*, 2014). With centriole formation stopped, a higher percentage of cells remains without centrioles/centrosomes at each round of cell division (**Fig. EV1A**). To avoid confusion, throughout this paper, centrosomes are defined as the PCM plus a parental centriole or a parental centriole including a pro-centriole. As previously reported (Tkach *et al*, 2022), we incubated MDA-MB-231 with DMSO (control) or 1000nM of CenB for 72 hours before subjecting them to immunofluorescence (IF) analysis (**Fig. EV1B**) by probing with the antibodies against the centriolar protein CEP192 and *γ*-tubulin (**Fig. 1A**). Centrosomes were identified as double-labelled puncta and manually counted per cell/nuclei. A relatively short CenB incubation of three days was chosen to obtain sufficient cells with full centrosomes removal, and simultaneously generate cells with a partial phenotype of 1 centrosome, as further internal control. Approximately half of the CenB-treated MDA-MB-231 population presented with zero centrosomes, while the remaining half still had one centrosome, and occasionally two (**Fig. 1B**). We then investigated whether the mesenchymal shape and size of both nuclei and cell soma changed in the absence of one or both centrosomes. We employed the generalist algorithm cellpose-SAM (Pachitariu *et al*, 2025) to segment cells, and quantified imaged features, alongside those of the segmented nuclei, with CellProfiler (CP) (Stirling *et al*, 2021) (**Fig. 1C**). An initial principal components analysis (PCA) of all extracted nuclei and cell soma features (e.g. area, solidity or convexity, bounding box, etc.) revealed no difference between the three groups of cells – containing two, one or zero centrosomes – and accordingly no clustering was present between controls and CenB-treated TNBC cells lacking centrosomes (**Fig. 1D-i, 1E-i**). For cells containing no centrosomes, only the nuclei shape, represented by the solidity/convexity value, was significantly different from that of controls. However, significance was minimal and could be explained by the aberrant division imposed by a spindle formed without centrosomes (**Fig. 1D-ii**). By contrast, cells that possessed one centrosome had a significantly increased size and significantly different shape in both nuclei and cells compared to controls (**Fig. 1D-ii, 1E-ii**). While the presence of a single centrosome seemed to alter the features of the TNBC cell, complete centrosome removal seemingly maintained the existing phenotype, suggesting negligible impact of centrosome function on already transitioned mesenchymal-like cancerous and metastatic cells.

**Figure 1.**
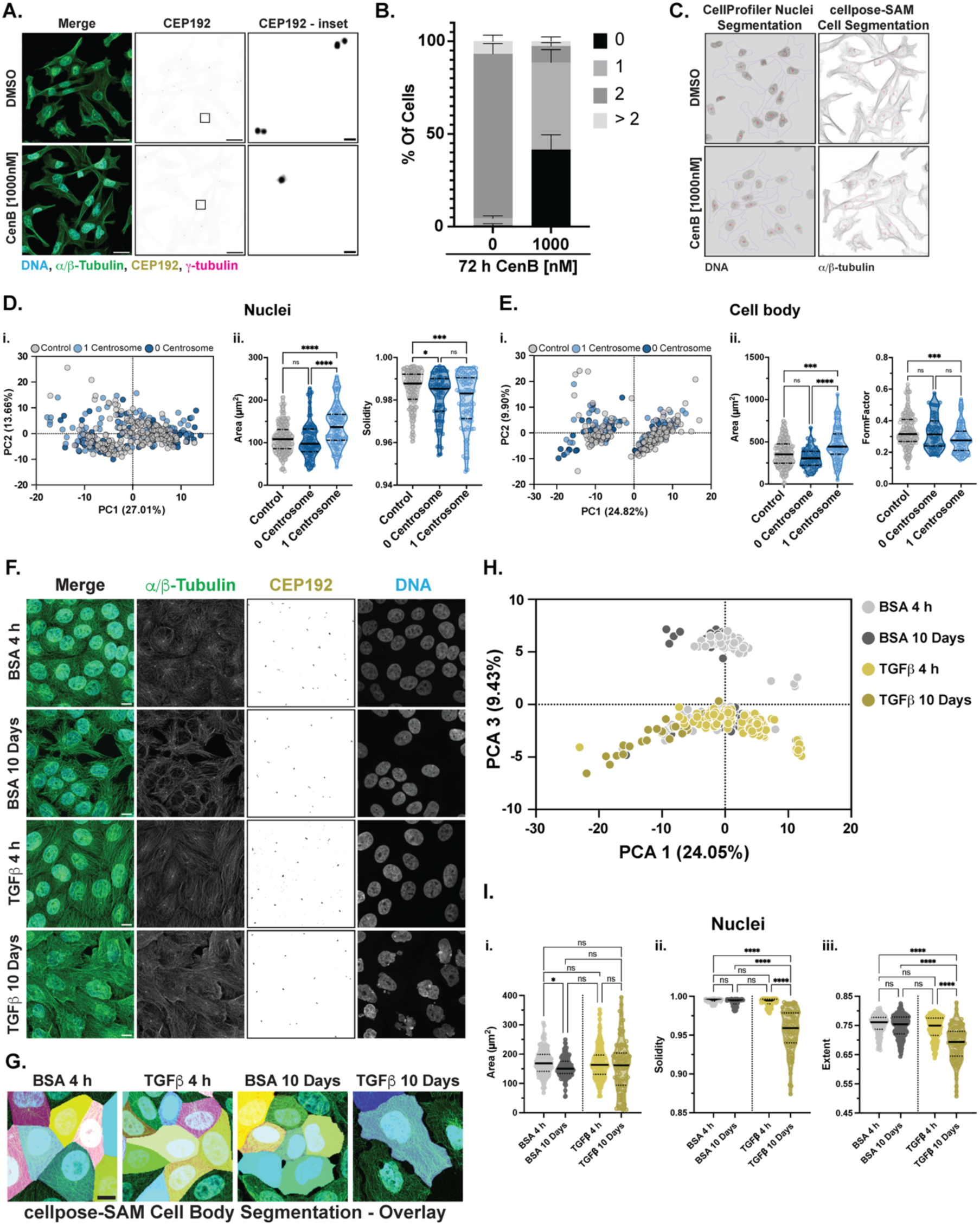
TNBC cells do not suffer changes in the absence of centrosomes, while MCF10A cells initiating the EMT re-organise their cytoskeletal network and alter nuclei shape. **A.** Representative AiryScan confocal IF images of methanol fixed MCF10A treated with DMSO or 1000nM CenB for 72 hours. Maximum intensity z-projections show merge of *α*/β-tubulin (green), CEP192 (yellow), γ-tubulin (magenta) and nuclei (DNA, Hoechst, cyan), and individual channel for CEP192 (grayscale, inverted). Scale bar: 10µm. Insert represents zoom-in of selected region (boxed) of CEP192 channel to better visualise centrosomes. Scale bars: 1µm. B. Bar plot shows quantification of centrosome number in MDA-MB-231 cells after 72h DMSO or 1000nM CenB treatment. Bars show mean ± SD of three independent experiments with a minimum of N=202 cells per condition. C. Representative images of Cellprofiler segmented nuclei (left panels) and cellpose-SAM segmented cell soma (right panels) for DMSO- or CenB-treated MDA-MB-231 cells. Outlines and ID object numbers of both nuclei and cell soma are overlaid on the inverted grayscale images of DNA and tubulin signals, respectively. D-E. MDA-MB-231 TNBC cells treated for 72 hours with either DMSO or CenB, and containing either 2 (control), 1 or 0 centrosomes. D-E i. PCA plot showing cells clustered according to all features extracted via CP analysis of cellpose-SAM segmented nuclei (left) and cell soma (right). Principal components (PC) 1 and 2 are plotted against each other; percentages show proportion of variance scores. D-ii. Violin plot quantification, showing median and interquartile range, of nuclei area (in µm^2^) and shape, defined by the solidity/convexity value (ObjectArea/ConvexHullArea). E-ii. Violin plot quantification, showing median and interquartile range, of cell soma area (in µm^2^) and shape, defined by the FormFactor value (4*π*Area/Perimeter^2^), of MDA-MB-231 treated with DMSO or CenB and containing either 2 (control), 1 or 0 centrosomes. In D-E, each data point in all graphs represents one nucleus or cell soma, derived from three independent experiments with a minimum N=86 for nuclei and N=58 for cell soma. Segmented cell bodies and nuclei touching the image border were excluded from analysis; similarly, cells containing 2+ centrosomes in the CenB-treated samples were removed from analysis. F. Representative AiryScan confocal IF images of 4% PFA-fixed MCF10A treated with TGFβ or BSA for 4 hours (4h) or 10 days. Maximum intensity z-projections show merge and individual channels of *α*/β-tubulin (green), CEP192 (yellow and inverted grayscale) and nuclei (DNA, Hoechst, cyan). Scale bar: 10µm. G. Representative masks obtained after cellpose-SAM segmentation and highlighting size differences in day 10 treated samples. Colored segmentation masks are overlaid on DNA/tubulin signals to show goodness of fit. H. PCA plot of DMSO- or TGFβ-treated cells at 4h or 10 days. Cells are clustered according to all features extracted via CP analysis of cellpose-SAM segmented cell somas. Principal components 1 and 3 are plotted against each other; percentage values show proportion of variance score. Each data point represents a single segmented cell, derived from two independent experiments with a minimum N=28. Segmented cells touching the border were excluded from analysis. I. Violin plot quantification, showing median and interquartile range, of i. nuclei area (in µm^2^) and shape, defined by ii. the solidity (ObjectArea/ConvexHullArea) and iii. extent values (ObjectArea/BoundingBoxArea). Each data point represents one nucleus, derived from two independent experiments with a minimum N=89. Segmented nuclei touching the border were excluded from analysis. For statistical significance, in all violin plots of Fig. 1D-E, except for the nuclei solidity, a ONE-way ANOVA followed by Kruskal-Wallis comparison test was performed. For the nuclear solidity, a ONE-way ANOVA followed by Tuckey’s multiple comparison test was used. For the violin plots of figure 1I, a two-way ANOVA followed by Tuckey’s multiple comparison test was performed. *p-values* meaning: non significant (ns)=> 0.05; *=0.05; ***=≤ 0.001; ****=≤ 0.0001.

### TGFβ-induced EMT in MCF10A cells promotes cytoskeletal rearrangement without impairing centriole number and ultrastructure

We next sought to understand which differences, if any, were imposed on the centrosome in response to the EMT. To study the EMT process from its inception, we induced it by stimulating the non-cancerous epithelial mammary cell line MCF10A with the cytokine TGFβ, or BSA as control (Zhang *et al*, 2014) (**Fig. EV1C**). We visualised the resulting transition via IF, by staining the cells with antibodies against **α**/*β*-tubulin, the centriolar protein CEP192 as well as a nuclear stain (DNA, Hoechst) (**Fig. 1F**). The same segmentation and feature extraction workflow as above was applied here (**Fig. 1G**). An initial PCA of all quantified features revealed a clear distinction between controls and TGFβ-treated cells, as well as a cluster only representing those cells which had been induced with TGFβ (dark yellow) for 10 days, pointing to significant and unique differences of cellular organisation imposed by the EMT initiation (**Fig. 1H**). Similarly as for MDA-MB-231 TNBC cells, alongside the changes in cellular structure, we also investigated the variations at the nuclear level. In the control setting at four hours (BSA 4 h) post-induction, nuclei were round, elliptical or ovaloid, and undented (**Fig. 1I**). Changes in nuclear size already appear after cellular aging (Pathak *et al*, 2021), as cells maintained in culture for 10 days (D10) demonstrated slightly lower area values (**Fig. 1I-i**). When supplemented with TGFβ, however, nuclei undergo a vast restructuring, resulting in nuclear shapes similar to those of bonafide cancerous cells (Zink *et al*, 2004; Verdone *et al*, 2015; Leggett *et al*, 2016a). Our results showed that, in MCF10A cells, already after a short burst of TGFβ, nuclei morphology varies, and this effect is accentuated over time after 10 days TGFβ incubation, with the appearance of micronuclei (visible as the percentage of cells with nuclei of small surface area in the lower quartile of the violin plot), and nuclei becoming irregular, elongated, or blebbing. These characteristics are captured by the lowering of the solidity/convexity measurement, which decreases from 1 (= perfect circle) to approx. 0.85, as well as a lower extent value (defined as area/bounding box area) (**Fig. 1I-ii,-iii**). Since the differences at the cellular and nuclear levels point to EMT initiation featuring vast internal rearrangements, we asked whether changes could also be visible at a numerical or ultrastructural level at centrioles. Via again IF analysis, we first quantified the variations in the number of centrioles present before and after TGFβ addition, but found no difference in the number of CEP192-positive puncta (**Fig. EV1D**). Next, we employed ultra-expansion microscopy (U-ExM) (Gambarotto *et al*, 2019) coupled with super-resolution imaging to visualise the ultrastructure of centrioles (Laporte et al,. 2024), by probing our samples with antibodies against acetylated tubulin - a localised, strong and reliable signal for centriole identification; and the resulting 3D centriole images were analysed with the segmentation software ilastik (Berg *et al*, 2019). Little difference was visible in the length of the centrioles between TGFβ-treated and control cells (**Fig. EV1E**): indeed, the small, but statistically significant, shorter centrioles at day 10 might be explained by an aging process in the cell culture system and possibly the experimental effects of cell overconfluence. Overall, with our protocol, we induced the EMT in non-carcinogenous mammary MCF10A cells and observed a drastic change in cell and nuclei morphology, but importantly, no difference in centriole number or ultrastructure.

### Lacking centrosomes significantly alters cellular and nuclear shape and size

As the MT network was instead heavily restructured in the presence of TGFβ, despite centrosomes appearing unaffected, we wondered what could be the effect if centrosomes, the main organisers of this network, were removed from cells, especially as more evidence in support of cancers presenting with centrosome-loss is emerging (Wang *et al*, 2020; Morretton *et al*, 2022b). To study the impact of the lack of centrosomes on the initiation of the EMT in breast tissue, we depleted centrosomes from the non-carcinogenous MCF10A cell line. Cells were treated for 72 hours with DMSO (control) or 1000nM CenB (**Fig. EV1F**), and subjected to the same IF analysis and quantification as TNBC cells, with centrosomes themselves identified by immunolabelling with anti-CEP192 and *γ*-tubulin antibodies (**Fig. EV1G**). After three days of CenB treatment, approximately 85% of cells lacked centrosomes (**Fig. EV1H**). We therefore proceeded to induce the EMT via TGFβ after centrosome removal, and investigated the effect of such centrosome depletion on the EMT process at day 0 (D0) and day 7 (D7) after TGFβ addition - hereafter referred to as before and after TGFβ (**Fig. 2A**). Unexpectedly, a first visualisation with phase contrast microscopy revealed that MCF10A lacking centrosomes, treated with both CenB and TGFβ (CenB+TGFβ), did not appear to have mesenchymal-like characteristic, similarly to cells treated with TGFβ still containing centrosomes (DMSO+TGFβ). Instead, TGFβ-treated cells lacking centrosomes (CenB+TGFβ) demonstrated ‘true’ epithelial characteristics, with tight and cohesive cobblestone-like morphology, strongly suggesting that a ‘classical’ EMT had not been initiated (**Fig. 2B**). To further analyze the underlying changes, we performed IF by staining MCF10A cells with antibodies against MTs (**α**/*β*-tubulin), actin (phalloidin), vimentin, and keratin14 (KRT14), as well as a nuclear marker (DNA, Hoechst) (**Fig. 2C**). IF analysis immediately confirmed the result of the phase contrast imaging (see **Fig. 2B**) and highlighted a dramatic change in shape, with narrower and elongated cells, as well as differences in cytoskeletal components, in the presence or absence of centrosomes during EMT initiation (**Fig. 2C, asterisks/arrowheads**). A Uniform Manifold Approximation and Projection (UMAP) analysis (McInnes *et al*, 2018), based on the resulting CP global measurements of cellpose-SAM segmented cells, illustrated that the treatment conditions form partially overlapping clusters. Untreated control cells containing centrosomes (DMSO+BSA D0 and D7) appear both as an overlap of each other and individually spaced, alongside centrosome-containing TGFβ-treated samples before induction (DMSO+TGFβ D0), where no change is also expected (**Fig. 2D**). CenB-treated cells lacking centrosomes (CenB+BSA D0 and D7) partially inhabit a region of the UMAP by themselves, highlighting a widespread impact of centrosomes removal that only marginally coincides with features promoted by TGFβ addition. Most importantly, TGFβ-induced cells (CenB+TGFβ and DMSO+TGFβ at D7) are clearly separated regardless of the absence or presence of centrosomes, underscoring significant differences imposed by TGFβ and supporting our finding that centrosome depletion impacts, and might be necessary, for EMT initiation. To more thoroughly unravel the changes between these conditions, we analysed independently the features that appeared most strikingly different, such as cell and nuclear size and shape (**Fig. 2E-F**). The spread of cell sizes was increased, albeit non-significantly, in centrosome depleted TGFβ-treated (CenB+TGFβ) cells before and after TGFβ addition. Treated cells at D7 (CenB+TGFβ), which lack centrosomes, were generally larger than matched CenB+BSA samples, potentially pointing to an effect on the cellular internal rearrangement, as well as more available surface for extending themselves since non-treated cells had a higher confluency (and less available space) after 10 days in culture (**Fig. 2E-i**). Importantly, by contrast, TGFβ-only-treated cells (DMSO+TGFβ) still containing centrosomes showed a much more protruded and elongated, spindle-like shape, as demonstrated by a low FormFactor value (calculated as 4*π*Area/Perimeter2), compared to all other conditions (**Fig. 2E-ii**). We then turned to discerning the changes imposed on the nuclei size and shape, as cancer is known to have a pleiotropic effect on this cell compartment (Zink *et al*, 2004; Comaills *et al*, 2016a) (**Fig. 2F**). Nuclei of CenB-treated cells lacking centrosomes, with or without TGFβ (CenB+BSA and CenB+TGFβ), were enlarged after treatment, possibly due to the misguided cell division occurring without a centrosome-orchestrated spindle pole (Hamzah *et al*, 2025) (**Fig. 2F-i**). Most notably, compared to untreated controls, nuclear roundness, measured by the solidity/convexity value, was significantly lower for centrosome lacking cells. Such decrease, while already present for centrosome-lacking samples before TGFβ treatment at D0 - at which point cells had already received 3 days of PLK4 inhibitor -, was further diminished after treatment at D7 for DMSO+TGFβ, CenB+BSA and CenB+TGFβ samples. This suggests that such changes are promoted both by the action of individual treatments, either the EMT initiation via TGFβ-induction or the lack of centrosomes, and are further exacerbated by their combination. To reinforce such statement: unlike for cellular shape, where morphological changes normally induced by TGFβ were prevented by centrosomes removal, indeed nuclei of TGFβ-induced cells lacking centrosomes (CenB+TGFβ) had the lowest mean solidity value, underlining a compound effect of both activated EMT and centrosome-dependent pathways that pushes the nucleus to an even more atypical form (**Fig. 2F-ii**). Crucially, to confirm that the visible effects on the EMT initiation were not just imposed by a slower cell cycle, we performed live time-lapse imaging of MCF10A cells treated with the various controls/conditions both at the starting point and after several days of TGFβ treatment (**Fig. EV2A-B**). We calculated the variation of the total number of nuclei present in the field of view over time as a proxy for cell proliferation (**Fig. EV2C**). MCF10A cells still proliferated in the presence of CenB (**Fig. EV2C**) and their wild-type p53 status did not trigger apoptosis, pointing instead to the failure of the surveillance mechanisms imposed by p16, which is found to be faulty in these cells and believed to be the driver of their spontaneous immortalisation (Cowell *et al*, 2005; LaPak & Burd, 2014; Qu *et al*, 2015). In the absence of centrosomes (+CenB samples), cell division is initially slowed, while the presence of TGFβ did not alter nuclei number compared to control. However, after CenB and/or TGFβ treatments at D7, there was no difference between the nuclei number in all conditions (**Fig. EV2C**). This suggests that while an initial hindering of the division rate might contribute to the delay in transitioning, it cannot fully explain how in the absence of centrosomes cells that continuously divide are maintained epithelial-like, suggesting instead that other factors/processes/pathways are most likely involved. Taken together these results suggest that while the change in cell shape normally resulting from the EMT induction is strongly reduced in absence of centrosomes, nuclear phenotypes following the EMT induction are actually exacerbated.

**Figure 2.**
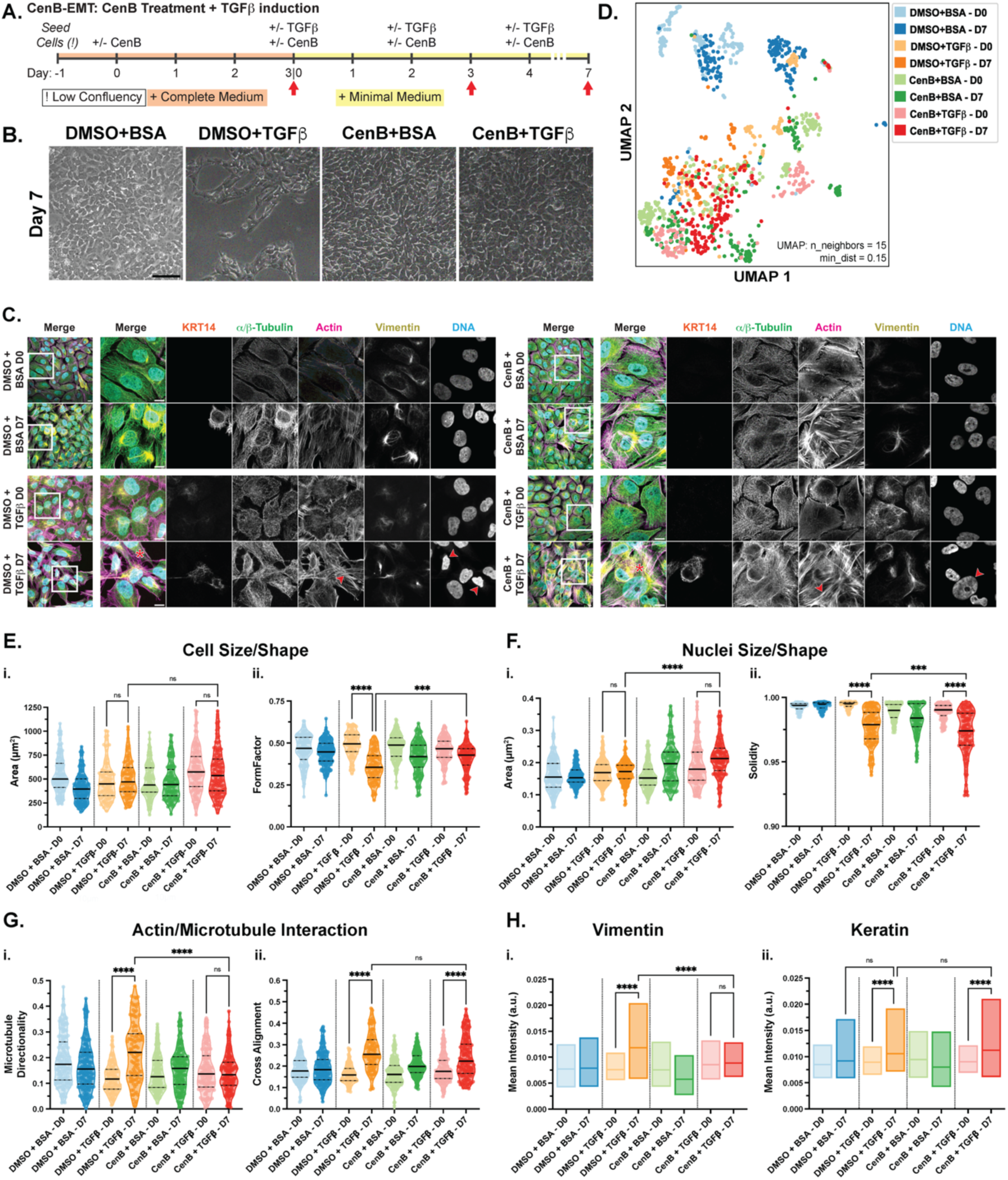
EMT induction in centrosome-depleted MCF10A cells promotes a change in cellular organisation. **A.** Schematic representation of CenB treatment followed by TGFβ induction protocol in MCF10A cells. Red arrows represent day/time of fixation (for IF) or harvesting (for WB and RNA-seq). **B.** Phase contrast images of MCF10A cells after TGFβ induction at day 7, with DMSO-BSA, DMSO-TGFβ, CenB-BSA or TGFβ-CenB. Scale bar: 100µm. **C.** Representative maximum intensity z-projections confocal IF images of 4% PFA-fixed MCF10A treated with a combination of CenB and TGFβ or controls at day 0 (D0) and day 7 (D7). First column for each condition panel shows the merge of the full field of view, while all other images are insets (boxed region) depicting the merge and individual channels of: KRT14 (orange), *α*/β-tubulin (green), actin (magenta), vimentin (yellow) and nuclei (DNA, Hoechst, cyan). Red asterisks at merge highlight the significant morphology change of DMSO-TGFβ and CenB-TGFβ-treated cells at D7; in the same samples, red arrowheads point to the actin stress fibres and irregular nuclei shape formed at D7. Scale bars: 10µm. **D.** UMAP plot of all CenB-TGFβ conditions and controls at D0 and D7. Each color coded data point clusters according to all features measurement extracted from CP analysis of cellpose-SAM segmented cells. Values of n_neighbors=15 and min_dist=0.15 are the hyperparameters inserted to visualise the clustering. **E.** Violin plot showing median and interquartile range of **i.** area (µm^2^) and **ii.** FormFactor (4*π*Area/Perimeter^2^) measurements of CenB-TGFβ-treated MCF10A cells soma for all conditions at D0 and D7. Measurements were obtained from CP analysis of cellpose-SAM segmented cells. **F.** Violin plot, showing median and interquartile range, of **i.** area (µm^2^) and **ii.** solidity (ObjectArea/ConvexHullArea) measurements of nuclei of MCF10A cells treated with CenB-TGFβ for all conditions at D0 and D7. Measurements were obtained from CP analysis of segmented nuclei. **G.** Graphical representation of differences in cytoskeletal components in MCF10A cells treated with CenB-TGFβ at D0 and D7. Structure tensor analysis measurements were obtained from CP-calculated features analysis of ilastik-segmented cytoskeletal components within cellpose-SAM segmented cells. Violin plot, showing mean and interquartile range, of **i.** microtubule directionality and **ii.** cross-alignment between actin and tubulin signal. **H.** Floating bar plot, showing min to max and line at mean values, of mean intensity values (in arbitrary units) of **i.** vimentin and **ii.** KRT14 signals. Throughout the figure, each data point represents one cell or one nucleus, and all datasets represented are derived from three independent repeats, with a minimum of N=98 per condition. Only cells or nuclei fully segmented within the field of view, and not touching the border, were considered for analysis. For statistical significance in all violin and bar plots an ordinary two-way ANOVA followed by Tukey’s multiple comparison test was performed. *p-values* meaning: non significant (ns)=> 0.05; ***=≤ 0.001; ****=≤ 0.0001. Only selected significance are shown; for a full description of significance see Expanded View Table Fig. 2.

### Lack of centrosomes alters cytoskeletal rearrangement imposed by EMT initiation

In order to understand the lack of shape change upon the EMT in absence of centrosomes, we further focused on specific features of the cytoskeletal components: actin, MTs (**α**/*β*-tubulin), vimentin and KRT14 (**Fig. 2G-H**). To better understand the rearrangement of the MT network nucleated in the absence of centrosomes, we calculated the order parameter of the **α**/*β*-tubulin signal and extracted the directionality of MTs. Cells containing centrosomes after TGFβ induction exhibited a higher directionality value compared to all other samples, which also demonstrated no change between each other. This suggests that, as expected, in the presence of centrosomes TGFβ promoted a cytoskeletal rearrangement typical of mesenchymal cells endowed with directed motion to invade surrounding tissues (Son & Moon, 2010) (**Fig. 2G-i**). As MTs and actin are known to interact extensively to promote cell structure and migration (Dogterom & Koenderink, 2019; Pimm & Henty-Ridilla, 2021; Nurmagambetova *et al*, 2023), we then focused on identifying the cross alignment of the signals between MTs and actin as a proxy for interaction of these two cytoskeletal components. Again after TGFβ treatment, TGFβ-treated cells containing centrosomes demonstrated a higher cross alignment value compared to all other samples except for CenB+TGFβ-treated cells lacking centrosomes (**Fig. 2G-ii**), the latter perhaps reflecting an altering of the EMT initiation where some components are primed for the transition while others are impeded, effectively limiting the EMT’s progression. To instead quantify the effect on the intermediate filaments vimentin and KRT14, we utilised the mean intensity signal per cell as a readout for protein amounts. Fluctuations in keratin levels as well as upregulation of vimentin are considered a hallmark of a mesenchymal state (Sharma et al, 2019). Indeed in accordance with previous literature (Chen *et al*, 2021), vimentin was significantly increased in TGFβ-treated cells containing centrosomes compared to all other conditions (**Fig. 2H-i**). The presence of KRT14 was instead a less clear EMT marker, presenting an increase in aged and TGFβ-treated cells, but also in samples lacking centrosomes treated with CenB (**Fig. 2H-ii**). Yet, such results potentially represent the natural state of MCF10A cells which already exhibit mixed luminal, basal and progenitor phenotypes (Chen *et al*, 2014; Qu *et al*, 2015), as well as KRT14 being a marker for singular cancerous cell rather than a collective feature (Cheung *et al*, 2016). To further confirm the presence of the EMT via its canonical markers, we performed a Western blot (WB) analysis, collecting samples both before and after TGFβ induction at day 0 and 7 (D0, D7) and at the halfway experimental point (D3). The key EMT-TF SNAI1 was visible only at D3 in the presence of TGFβ, regardless of centrosome content, demonstrating its expected activation promoted by TGFβ as a driver of the epithelial-to-mesenchymal switch (**Fig. EV2D**). Fibronectin and N-cadherin were both present at D7 in TGFβ-treated samples, but not in untreated controls or centrosome-lacking CenB-only-treated cells (**Fig. EV2D**). Overall, the WB analysis confirmed the happening of the EMT via the upregulation of classical markers such as fibronectin, N-cadherin and SNAI1, in TGFβ-treated samples, regardless of centrosome presence or absence. In CenB-treated samples lacking centrosomes KRT14 is also increased, while vimentin is not (see **Fig. 2C-2H**), as instead expected for cells undergoing the EMT - and visible in the WB. Similarly, MTs are highly directional in the presence of centrosomes and TGFβ compared to all other samples, yet the alignment between its cytoskeletal components is equivalently increased in the EMT initiated cells regardless of centrosomes content (see **Fig. 2G**). Furthermore, while the area of cells lacking centrosomes is greater, their shape is less fusiform than that of TGFβ-treated cells containing centrosomes (see **Fig. 2E**). It has been demonstrated that cells lacking centrosomes rely on alternative MTOCs, such as the nuclear membrane, cell cortex and Golgi apparatus (Sanchez & Feldman, 2017; Akhmanova & Kapitein, 2022; Van Grinsven & Akhmanova, 2025). We therefore examined the Golgi apparatus to establish if any difference was present at this key organelle. After analysing the Golgi-specific marker GM130 via IF (**Fig. EV2E**) in our CenB-TGFβ treated MCF10A, we found that although the lack of centrosomes slightly enlarged Golgi size (bounding box area) at D7, it did not prevent TGFβ-induced Golgi disorganization and fragmentation (number of children) observed during the EMT and in cancer (Bui *et al*, 2021), indicating that some EMT-associated changes may still proceed despite only partial changes in cellular morphology (**Fig. EV2F**). Such results further underline an alteration or limit in the initiation of the EMT when centrosomes are lacking. All together the results obtained from the morphological, distribution and quantification analyses highlight that, in the absence of centrosomes, the EMT is not occurring as expected, in a linear or classical manner, and that cytoskeletal rearrangements are not completely happening.

### RNA-seq reveals a misregulation of EMT hallmarks and ECM remodelling in response to centrosome removal

Our next question therefore was to ask what changes occur at the RNA level and what genes were being activated or repressed to potentially explain the cellular variations seen during the EMT initiation in centrosome-lacking cells. By performing RNA-sequencing, our next aim was to obtain an overview of the transcription pathways being affected by EMT induction in the presence and absence of centrosomes. Initial exploratory quality control of our triplicate samples showed good correlation/distribution between the conditions (**Fig. 3A**, **Fig. EV3A**). The overall top 50 differentially expressed genes already showed expected variations for samples treated with TGFβ, such as the decrease of S100A9 and S100A8, calcium binding proteins well known to be cancer effectors (Salama *et al*, 2008), as well as the increase in metalloprotease (MMP2/MMP10/ADAM19) and fibronectin (FN1), both classic EMT markers (**Fig. EV3B**). A 3D PCA plot further highlighted the different gene expression programs. Cells at D0 were clearly separated between cells containing or lacking centrosomes but still occupied the same region of the PCA. After the different treatments all conditions moved from the initial region but all in different directions, representing significant transcription changes and differences imposed by the centrosomes during EMT initiation (**Fig. 3A**). In accordance with the PCA results, the presence of TGFβ alone induced an upregulation and downregulation of more than 3000 and 2000 genes respectively (**Fig. 3B-i**), while comparing the TGFβ-induced genes in presence and absence of centrosomes showed a difference of regulation in excess of 100 upregulated and 150 downregulated genes (**Fig. 3B ii**). To confirm that the affected genes included those specifically regulated by TGFβ, we compared the expression levels of the genes upregulated (TGFβ-induced) and downregulated (TGFβ-repressed) by TGFβ in presence and absence of centrosomes (**Fig 3C**). The results showed a significant reduction of TGFb-induced gene regulation in absence of centrosomes. As depicted in the heatmap Fig 3D, this effect is not only observed for genes relating to TGFβ signalling but also for ECM remodelling and overall EMT hallmark regulated genes, that get less activated in the absence of centrosomes in CenB+TGFβ treated cells compared to TGFβ-induced cells after treatment (**Fig. 3D**, **EV3C-E**). Further genes, upregulated by TGFβ in the presence of centrosomes, were not activated in their absence, suggesting a necessary function of this organelle in promoting a classical EMT initiation (ex. LRRC15, COL3A1, SLIT2, etc.) (**Fig. 3B, 3D, EV3E**). Collagen genes upregulated in the TGFβ-only condition, such as COL4A1, COL4A4, COL6A3, COL5A3, COL8A2, and COL12A1, were less activated in CenB+TGFβ samples lacking centrosomes (**Fig. EV3D,** see Material and Methods: Data Availability). Similarly, key MMPs upregulated in TGFβ-treated cells (MMP7, MMP9, MMP10) were less activated in the presence of TGFβ and CenB (CenB+TGFβ); other genes involved in cancer biology (ex. RAC2, CLU, ADAMTS5) as well as some key classical EMT transcription factors (ZEB1 and SNAI1) were also not activated, further highlighting the modulatory effect of centrosome depletion on the EMT already from transcription (**Fig. 3B, 3D,** see Material and Methods: Data Availability). Remodelling of the ECM is a well-known feature of the EMT and a key process of the metastatic program (Chen *et al*, 2025), involving alterations in cell-to-cell and cell-to-ECM communication, adhesion and signalling, which ultimately reinforce the EMT. Paramount to the transition are also the links between the cell membrane and its junctions, and the cytoskeletal elements responsible for structure, transport, migration and signals between intra and extracellular inputs bidirectionally (Janiszewska *et al*, 2020). Similarly to the transcription factors, MMPs and collagens, some members of the cell-cell and cell-matrix communication pathways were inversely regulated in TGFβ-treated cells in the presence and absence of centrosomes, such as ITGA4, CDH5, ANK1, MYLK, etc (see Material and Methods: Data Availability). Overall the RNA-sequencing analysis confirmed that the lack of centrosomes has a partial effect on the EMT canonical transcriptional program induced by TGFβ, mitigating the upregulation of some gene networks including ECM remodelling and cell adhesion/migration related genes.

**Figure 3.**
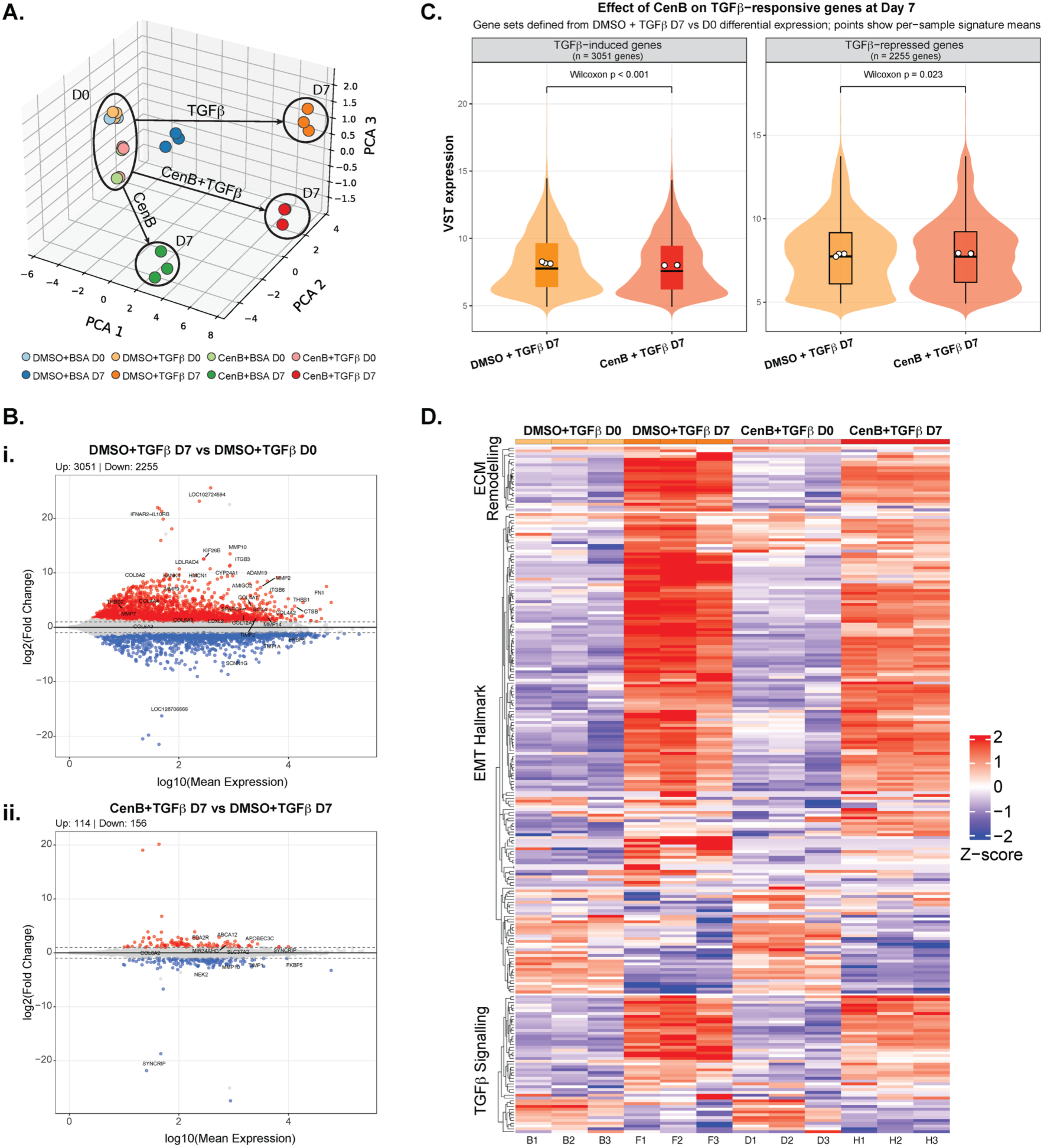
RNA-seq highlights differences in cytoskeletal and ECM pathway regulation in the absence of centrosomes. **A.** 3D PCA plot of RNA-seq samples of MCF10A cells treated with CenB-TGFβ at D0 and D7. Color coded data points represent individual RNA extraction repeats and clusters according to condition and days of treatment. Circles and arrows highlight cluster changes between D0 and D7 in the different conditions. **B.** MA plots of differential gene expression (DGE) between RNA of MCF10A cells treated with **i.** DMSO-TGFβ at D7 vs D0 and **ii.** CenB-TGFβ vs DMSO-TGFβ at D7. Red (above 0) and blue (below 0) dots represent upregulated and downregulated genes, respectively, in the first condition relative to the reference condition (i.e., D7 relative to D0 in the left panel and CenB relative to DMSO at D7 in the right panel). Gene names of particular interest and opposing effects are shown on graphs. **C.** Violin plot representing the effects of CenB on TGFβ-responsive genes after TGFβ treatment. Variance Stabilising Transformation (VST) expression levels are illustrated for upregulated (TGFβ-induced genes, left panel) vs downregulated (TGFβ-repressed, right panel) genes. For significance a Wilcoxon rank-sum test was performed and the *p-value* obtained specified. **D**. Heatmap of DGE for genes involved in ECM remodelling, EMT hallmark and TGFβ signalling in TGFβ only and CenB-TGFβ conditions, before and after treatment. For a full list of gene names in all tested conditions refer to heatmap on EV3D and Material and Methods: *RNA sequencing Differential Expression Analysis (DEA)*.

### EMT-induced paxillin relocalisation at focal adhesions does not occur in the absence of centrosomes

We hypothesized that centrosome loss could impair the coupling between the actin cytoskeleton and cell-ECM adhesions during TGFβ-induced EMT. To test this, we focused on paxillin, a key regulator of focal adhesion dynamics linking integrin signalling to actin remodelling (Turner et al, 1990; Deakin & Turner, 2011). Although paxillin itself was not differentially expressed in our RNA-seq dataset, our hypothesis emerged from integrating its established biological function with the altered expression of several paxillin-associated regulators, including the upregulation of PAK2, FAK and β-catenin (Malla et al, 2025) and the downregulation of Kindlin-1 (Li et al, 2025), irrespective of centrosome status. Thus, rather than predicting changes in paxillin expression, we reasoned that perturbation of its regulatory network would instead alter its organisation during the EMT. Given the established roles of paxillin in cell migration, cell-matrix communication and breast cancer progression (Malla et al, 2025; Shortrede et al, 2016; Xu et al, 2022; Liu et al, 2023), we investigated whether centrosome depletion affects paxillin organisation during the EMT. We again performed IF on the CenB-TGFβ-stimulated MCF10A before and after treatment and immunolabelled our samples for paxillin, actin (via phalloidin staining), as well as **α**/*β*-tubulin and a nuclear stain (DNA, Hoechst). In control settings, paxillin was distributed throughout the cell, perinuclearly and within the nucleus, and found as puncta at the tip of actin fibers or at the cell’s periphery, expectedly (**Fig. 4A**). We used two different complementary coefficients, Manders’ and Pearson’s, (Manders *et al*, 1993; Dunn *et al*, 2011) to quantify the mutual relationship between the paxillin and actin signal after the different treatments. The Manders’ score, that reports on signal fraction overlap, clearly identified a proportion of paxillin overlapping with actin that is much greater in TGFβ-only-stimulated only in centrosome-containing sample, while all other conditions show similar and non-statistically significant values (**Fig. 4B-i**). Similarly, the results of Pearson’s correlation, which measures the relationship between the variation in concentration of the two molecules, indicated that only centrosome-containing TGFβ-induced samples correlated less negatively than all other samples, pointing to less variation in the concentration of paxillin and actin within cells (**Fig. 4B-ii**). Both results strongly support the notion that only centrosome-containing TGFβ-treated cells showed increased localisation of paxillin at focal adhesions suggesting a necessary role of centrosomes in the relocalisation of paxillin in the EMT.

**Figure 4.**
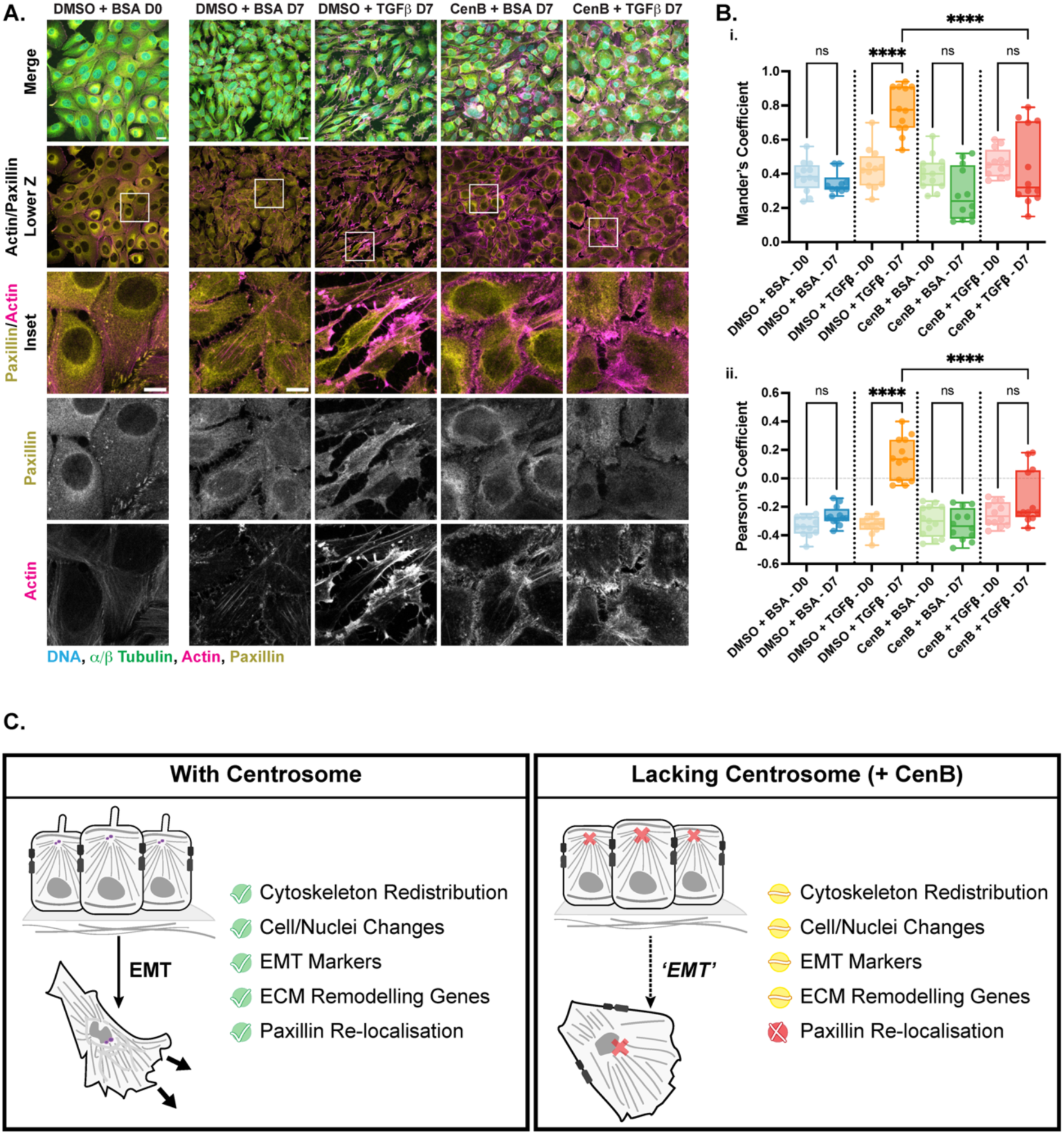
Paxillin/actin colocalisation at focal adhesions is lessened in EMT-induced MCF10A devoid of centrosomes. **A.** Representative confocal IF images of 4% PFA-fixed MCF10A treated with a combination of CenB and TGFβ or controls at day 0 (D0) and day 7 (D7). Maximum intensity z-projections (MIP) show merge of paxillin (yellow), actin (phalloidin, magenta), *α*/β-tubulin (green) and nuclear stain (DNA, Hoechst, cyan). Lower Z row shows MIP of a stack of only four slices: one below the coverslip and three at/above, to capture focal adhesions only. Scale bar: 20µm. Paxillin/Actin insets (boxed region) show zoom-in of IF images for the merged and single grayscale channels. Inset scale bar: 10µm. **B.** Box and whiskers graph showing, min to max with line at median, of **i.** the Manders’colocalisation coefficient and **ii.** Pearson’s correlation coefficient between paxillin and actin signals. For both Manders’ and Pearson’s coefficients the value above autothreshold of CH1 (actin) were taken. Throughout the figure, each data point represents one image frame containing multiple cells, and all datasets represented are derived from three independent repeats and with a minimum of N=12 fields of view per condition. For statistical significance in all box plots an ordinary two-way ANOVA followed by Tukey’s multiple comparison post-hoc test was performed. *p-values* meaning: non significant (ns)=> 0.05; ****=≤ 0.0001. Only selected significance are shown, for a full description of significance see Expanded View Table Fig. 4. **C.** Schematic representation of the effects of lacking centrosomes in the EMT and cancer initiation. In the presence of centrosomes, TGFβ-stimulated cells initiate and undergo the EMT: cells change their morphology to a spindle-like appearance, with shifts in nuclear and cell body shape and size, upregulate classical EMT markers, upregulate ECM remodelling program and localise paxillin to the focal adhesions potentially favouring an invasive behaviour. After centrosome removal, TGFβ-stimulated cells do not acquire a fully mesenchymal morphology (Cytoskeleton Redistribution and Cell/Nuclei changes), suppress the classical EMT markers, reduce ECM remodelling genes, and prevent cytoskeletal paxillin re-localisation, overall promoting a delay, hampering or alternative EMT state.

## Discussion

The EMT process underpins cancer aetiology and is a key driver of metastatic potential, the responsible cause for high mortality in BC patients. Cells undergoing the EMT extensively transform their cellular structures: from their nuclei to their cytoskeleton, their signalling pathways and extracellular environment. The role of centrosomes in cancer has been long postulated and evidence of their key involvement abounds, yet their contribution to the EMT programme and its initiation is not fully understood. This study aims to identify the importance and actions of centrosomes in the transition of human mammary cells from an epithelial to a mesenchymal phenotype. From our results, initiation of the EMT in non-carcinogenic mammary cells rearranged their cytoskeleton and reshaped their nuclei, supporting current literature (Comaills *et al*, 2016b; Leggett *et al*, 2016b; Garcia-Murillas *et al*, 2019), without any impact on centrosomes number or ultrastructure. In the absence of centrosomes, however, such morphological restructuring was lost, and transitioning cells presented with a similar cytoskeleton to that of epithelial cells. Nuclei shape, however, still became atypical and classical EMT markers were being expressed, pointing to a limited EMT: a delay or the activation of an alternate pathway in its progression. Analysis of the RNA transcriptome further revealed that centrosome depleted cells prevented the ECM remodelling gene expression from unequivocally resembling that of a cancerous phenotype, as well as stabilising the presence of key cell adhesion molecules, further impeding the transition from an epithelial towards a quasi-mesenchymal state. Based on these effects, an investigation on the centrosome’s role in adhesion molecule relocalisation found that, in absence of centrosome, paxillin did not relocalise upon EMT induction. Taken together, these results indicate that centrosomes are required for a canonical EMT transition (**Fig. 4C**). Conversely, already mesenchymal-like cells do not suffer from the loss of centrosomes, and whatever impact such, complete or partial, loss has on metastatic cells is still to be uncovered.

Many questions have emerged from our investigation, and further studies are required to fully unravel the impact of centrosomes on the initiation of the EMT in mammary cells. For examples, in regards to nuclear shape changes, it will be paramount to separate and sequence the transcriptome of exclusively amorphous nuclei seen in the absence of centrosomes from those with normal shape, to highlight which differentially regulated pathways or genes reinforce or promote this phenotype (Cosenza *et al*, 2025). It would be of interest to replicate the above experiments in the presence of the ECM and within a 3D structure, to evaluate the contributing effect of the chemical remodelling and physical forces of the matrix as well as polarising forces of 3D architecture, especially as compositional ECM changes - loss of biochemical and mechanical homeostasis - are known to contribute to cancer (Higgins *et al*, 2025). Such experiments would also allow the monitoring of the invasive behaviour of cells, as either clusters or single cells, and how they rearrange their cytoskeleton to promote migration in a 3D tumour microenvironment. Indeed, in this context, a closer inspection on paxillin, its physical interaction with integrins, other adaptor proteins such as vinculin and actin, and its role in triggering RhoGTPas as well as its influence on migration, might reveal mechanistic insight into the pathways that are directly coordinated by centrosomes. Understanding its trafficking towards the plasma membrane, via a deregulated MT network, might also explain why we recorded a decrease in its co-occurrence and correlation at the focal adhesion in cells lacking centrosomes. Having deduced the involvement of paxillin from the differentially expressed ECM and adhesion genes obtained from the transcriptome analysis of treated MCF10A lacking centrosomes, a detailed RNA sequencing of TNBC MDA-MB-231 is also highly anticipated. The differential gene expression of TNBC cells containing one or zero centrosomes would reveal on one side the pathways that are directly targeted and modulated by the CenB without the interfering effects to TGFβ signalling, and on the other, importantly, the singular contributions of centrosome loss to the mesenchymal state.

Our study further highlights a role for centrosomes themselves, either passive or active, in cancer establishment that is unrelated to supernumerary and structural centrosome aberrations. The pathogenic mode of action of centrosomes amplification includes disruption of the tissue architecture, via increased MT nucleation and Rac1 signalling (Lingle & Salisbury, 1999; Godinho *et al*, 2014); increase in reactive oxygen species that promote secretion of pro-invasive factors as well as extracellular vesicles and exosomes, that alter the tumor microenvironment and exacerbates cancer features (Arnandis *et al*, 2018; Adams *et al*, 2021). Conversely to the detailed literature on centrosome amplification, centrosome loss is a feature of cancers that has only recently been detected and is slowly being investigated *in cell*. CenB treatment produced acentrosomal cells that still formed functional spindles, often constructed by PCM proteins, that coordinated a slower, more error-prone cell division (Watanabe *et al*, 2020). Centrosome removal in non-carcinogenic prostate epithelia spurred aneuploid and multinucleated cells capable of generating malignant tumors when xenographed in mice, possibly by introducing chromosomal instability (Wang *et al*, 2020; Yang *et al*, 2026). In an organoid model of colorectal cancer, centrosome loss repressed growth and survival in normal tissue but mediated growth defects in patient-derived samples (Bourmoum *et al*, 2024). Indeed centrosome loss, and ensuing prolonged mitosis, triggers the USP28-53BP1-p53-p21 signaling pathway to function as a mitotic surveillance system and prevent unfit daughter cells with mitotic errors from proliferating (Lambrus *et al*, 2016). While most studies described here focus on the cellular response to missing centrosomes, our investigation targets the impact of centrosomes at the stages of cancer initiation, when cells turn from epithelial to mesenchymal. When absent at the EMT initiation, centrosomes prevent a reorganisation of the cytoskeleton and reduce the expression of some genes critical for ECM, which results in cell adhesion and migration proteins, such as paxillin, potentially impacting motion and cell-cell communication.

Significantly, the effects of induced centrosome removal in cultured cell lines might not fully reflect what occurs within tumoral tissues; and while - to our knowledge - not yet described, centrosome loss might well be a feature of BC, as for prostate and ovarian cancers (Morretton *et al*, 2022b; Yang *et al*, 2026). Centrioles/centrosomes are also the building blocks upon which cilia form. Primary cilia are fundamental signalling hubs found to be both overactive as well as completely absent in cancers in a context-dependent manner (Collinson & Tanos, 2025). We cannot exclude that by removing centrioles/centrosomes, and thus effectively removing cilia, the hindering/delaying effects of the EMT evident on the BC cells are the result of misregulated or absent cilia signalling (Wilson *et al*, 2021), and if/to what extent this might be the case needs further investigation.

As PLK4 inhibitors are progressing through clinical trials, parallel scientific discoveries are increasing the knowledge to maximise their inhibitory sensitivity: by, for example, targeting cancers with higher levels of TRIM37, increased β-catenin signalling or the cluster supranumerary centrosomes, or by employing them in combination with other drugs (Soria-Bretones *et al*, 2026). Our study additionally highlights the importance of understanding centrosome removal at different stages of cancer progression, and their fundamental yet different effect in still epithelia and/or already mesenchymal cells. Indeed, centrosomes appear to possess a double-edge effect during the EMT: capable of promoting the transition when cells are still epithelial, but seemingly less consequential when the transition has already occurred. As more and more therapeutic strategies hinge on modifying centrosomes and/or their related pathways, this work cautiously underscores the possible limitations of the impact or efficacy of centrosome inhibitors on already cancerous cells, and highlights the importance of drug delivery timing, at the cusp of the EMT.

## Materials and Methods

### Methods

#### Cell Culture

MCF10A cells were obtained from Cell Line Service (CLS) (lot number 305026-250123SF) and cultured according to standard protocol (Debnath *et al*, 2003) in Dulbecco’s Modified Eagle Medium: Nutrient Mixture F-12 (D-MEM/F-12) (Gibco, 12634010), and further containing 5% horse serum (Gibco, 26050088), 20 ng/ml Epidermal Growth Factor (PeproTech, AF-100-15), 0.5 mg/ml hydrocortisone (SigmaAldrich, H0135-1MG), 10 μg/ml insulin (SigmaAldrich, I1882), 100 ng/ml cholera toxin (SigmaAldrich, C8052) and 1% Pen/Strep (SigmaAldrich, P0781). Triple negative breast cancer cell line MDA-MB-231 was maintained in DMEM High Glucose with L-glutamine and phenol red (Gibco, Cat#: 41965062) supplemented with 10% FBS (Gibco, A5256501) and 1% Pen/Strep (SigmaAldrich, P0781). Cells were maintained in a 5% CO_2_-humidified incubator at 37°C, and grown for 4 to 5 weeks in fresh media and discarded above passage 24. Cell authenticity and health was checked by STR profiling and routinely tested negative for mycoplasma contamination by PCR.

#### TGFβ induced transformation and immunofluorescence

A variation of the protocol from Zhang et al. (Zhang *et al*, 2014) was used to induce the EMT in MCF10A cells via the addition of recombinant human TGF-β1 (PeproTech, 100-21C). Cells were plated on clean high-precision 12mm cover slips (ThorLabs, CG15NH) in complete medium.Once cells were 80% confluent, media was replaced with pre-warmed minimal media, composed of DMEM/F-12 containing only 2% horse serum and 100 ng/ml cholera toxin - and excluding all other components. Cholera toxin was kept in the minimal medium to prevent spontaneous elongated, mesenchyme-like phenotype. The minimal media contained either 5 ng/ml TGF-β1 or 0.1% BSA as vehicle. Cells were allowed to transform undisturbed at 37°C and 5% CO_2_ for up to 10 days, and minimal medium was replaced every 48h. Either at 4h, 6 or 10 days after TGFβ addition, cells were washed once in 1xPBS and fixed in 4% PFA in 1xPBS (ThermoFisher, 416780250) for 20min at RT; cells were then either processed immediately for immunofluorescence or stored at 4°C in 1xPBS with 0.3% NaN_3_ (SigmaAldrich, S2002). For IF, after fixation cells were permeabilised in 1xPBS + 0.05% Triton-X (SigmaAldrich, X100) for 15min at RT, followed by two washes in 1xPBS for 5min each. Cells were then blocked in 3% Bovine Serum Albumin (BSA) (Cell Signalling, 9998S) in 1xPBS for 45min at RT, followed again by two washes in 1xPBS for 5min each. Primary antibodies were diluted in 1% BSA in 0.025% PBS-Tween20 (PBS-T) (Promega, H5151) and incubated on the sample for 2h at RT. Samples were then washed three times for 10min each at RT with 0.1% PBS-T and further incubated in the dark with secondary antibodies diluted in 1% BSA in 0.025% PBS-T for 1h at RT. A further three washes of 10min each with 0.1% PBS-T followed, before cells were mounted on microscope glass slides (Carl Roth, H870.1) with the aid of ProLong Gold antifade mountant (ThermoFisher, P36930). Samples were left to solidify on the slides overnight in the dark before imaging. Cells were either probed with antibodies against centriolar and cytoskeletal components or for EMT markers. Primary and secondary antibodies used for this IF were: anti-vimentin (1:2000, Proteintech, 10366-1-AP), anti-centrin (1:1000, SigmaAldrich, 04-1624), anti-**α** and anti-*β* tubulin (1:3000, ABCD antibodies, ABCD_AA345 and ABCD_AA344, respectively), donkey anti-mouse Alexa Fluor 488 (1:10000, ThermoFisher, A-31571), goat anti-guinea pig Alexa Fluor 568 (1:10000, ThermoFischer, A-11075), donkey anti-rabbit Alexa Fluor 647 (1:10000, ThermoFisher, A-31573), and goat anti-rat Alexa Fluor 647 (1:10000, ThermoFischer, A-21247). Cell nuclei were counterstained with Hoechst (1 µg/ml, Invitrogen, H1399).

#### Ultra-expansion microscopy of MCF10A treated cells

ExM was performed following the ultra-structural expansion microscopy protocol by Gambarotto et al. (Gambarotto *et al*, 2019). Briefly, 4% PFA or methanol-fixed cover slips samples were cross-linked by incubation for 3.5h at 37°C with a solution containing 2% acrylamide (AA 40%, SigmaAldrich, A4058) and 1.4% formaldehyde (FA 36.5-38%, SigmaAldrich, F8775) in 1x PBS. Cover slips were then placed on a drop of chilled monomer solution - composed of 19% (wt/wt) sodium acrylate (SA, Merck, 408220), 10% (wt/wt) acrylamide, 0.1% (wt/wt) N,N’-methylenbisacrylamide (BIS, Merck, M7279) in 1xPBS - and supplemented with 0.5% (wt/wt) TEMED (ThermoScientific, 17919) and 0.5% (wt/wt) APS (ThermoFisher, 17874), with the cells facing the monomer solution. Samples were transferred to a pre-chilled humid chamber and left undisturbed at 37°C for 1h. After gelation, gel samples were gently tipped from the cover slips into denaturation buffer - 200 mM SDS (Serva, 20765.01), 200 mM NaCl and 50 mM Tris (SigmaAldrich, T1503) in distilled H_2_O at pH 9.0 - and left to detach by agitation at RT for approximately 30min. Detached gels were placed in 2ml Eppendorf tubes and submerged in denaturation buffer; tubes were placed to shake vigorously at 450RPM at 95°C in a thermomixer for 1.5h. After denaturation, gels were placed in copious amounts of distilled water and allowed to expand for 1h at RT, changing the water once in between. Gel size was measured with the aid of a calliper. An expansion factor of 4.00 was recorded on average. Gels were then shrunk by adding 1xPBS twice for 15min and then blocked for 30min at 37°C in 3% BSA and 0.5% Triton-X in 1xPBS. Excess BSA solution was carefully removed and gels were washed with 1x 0.1% PBS-T for 10min. Gels were then incubated with primary antibodies diluted in 1% BSA in 0.05% PBS-T overnight shaking at 4°C. The next day, gels were washed 3x with 1x 0.1% PBS-T for 10min each and then placed in the dark shaking for 2h at 37°C within the secondary antibody solution diluted also in 1% BSA in 0.05% PBS-T. Gels were then washed 3x with 1x 0.1% PBS-T for 10min each and incubated for 10min in a nuclear stain solution containing 1 μg/ml Hoechst (Invitrogen, H1399) in 1xPBS. Another wash in 1x PBS for 10min followed, before finally placing the gels in distilled water overnight to allow for a second round of expansion. Gels were ready to be imaged by mounting on lightly covered Poly-L-Lysine (SigmaAldrich, P8920) coated high-precision glass coverslips, to prevent drifts. Primary and secondary antibodies used for centrosome staining in U-ExM are Acetyl-**α** Tubulin (Lys40) (1:500, Invitrogen, 32-2700) and donkey anti-mouse Alexa Fluor 488 (1:10000, ThermoFischer, A-21202), respectively.

#### Centrinone B treatment and immunofluorescence

Experiments were performed based on the originally described Centrinone B (CenB) protocol (Wong *et al*, 2015). MCF10A or MDA-MB-231 were cultured in their complete medium respectively as described above. CenB powder (TOCRIS, 5690) was diluted in DMSO to a stock concentration of 1mM according to manufacturer’s instructions and kept in aliquots at - 20°C. An aliquot of CenB was thawed freshly for each experiment and used once only. Cells were seeded on cleaned 12mm high-precision cover slips to a low confluency of 20%. The next morning, media was replaced with pre-warmed fresh media containing either 1000nM CenB or DMSO (as vehicle) - during which time, the PLK4 inhibitor would have acted to prevent centriole formation resulting in effective centrosome loss. After 72h, cells were washed with 1xPBS and fixed in ice cold methanol (Avantor, 8402) at -20°C for 7min. Cells were washed twice in 1xPBS and either stored at 4°C in 1xPBS with 0.3% NaN_3_ or used directly for IF, which was performed as described above. Primary and secondary antibodies utilised for this IF were: anti-CEP192 (Proteintech, 28700-1-AP), anti-*γ*-tubulin (Abcam, ab27074), anti-**α** and anti-*β* tubulin (ABCD antibodies, ABCD_AA345 and ABCD_AA344 respectively), donkey anti-mouse Alexa Fluor 488 (ThermoFischer, A-21202), goat anti-guinea pig Alexa Fluor 568 (ThermoFischer, A-11075), and donkey anti-rabbit Alexa Fluor 647 (ThermoFisher, A-31573). Cell nuclei were counterstained with Hoechst (1 µg/ml, Invitrogen, H1399).

#### CenB-TGFβ treatment, immunofluorescence and western blotting

Cells were seeded on clean high-precision 12mm cover slips at a low 20% confluency, in complete medium and allowed to attach overnight. The next day, the medium was replaced with fresh media containing either 1000nM CenB or DMSO as vehicle. Cells were then incubated for 72h unperturbed at 37°C and 5% CO_2_. After three days, cells were switched to minimal media (2% horse serum plus 100ng/ml cholera toxin) and further stimulated with either TGFβ or 1% BSAl (as vehicle); importantly, cells that had received CenB were again supplemented with CenB, to maintain centrosome ‘loss’ and prevent centrosome overduplication. Again, cholera toxin was kept in the minimal medium to prevent spontaneous elongated, mesenchyme-like phenotype. In total, four conditions — each containing a treatment or control — were established for comparison: DMSO+BSA, as control for both chemicals added; DMSO+TGFβ, as control for the EMT; CenB+BSA, as control for the effects of centrosome loss; and CenB+TGFβ as the treatment of interest. Fresh medium containing both CenB and TGFβ treatments, in their respective combination and conditions, was replaced after 48h twice more, on day 2 and day 4 after the initial TGF*β* stimulation. Cells were fixed on day 0 and day 7 after TGFβ induction (intended respectively as day 4 and day 10 after seeding and after three days of CenB treatment). At the correct timepoints, cells were quickly rinsed in 1xPBS and fixed in 4% PFA in 1xPBS for 20min at RT, and 10min at RT for paxillin experiments. Fixed cells were rinsed twice again after fixation and then stored at 4°C in 1xPBS with 0.3% NaN_3_ or immediately processed for IF. IF was performed as described above, with the only variation that the incubation time of the primary antibody was extended to 3 hours at RT, instead of 2 hours. CenB-TGF*β* treated cells were stained with the following primary antibodies: anti-**α**/*β* tubulin (1:3000, ABCD antibodies, ABCD_AA345 and ABCD_AA344, respectively), anti-GM130 (1:3000, Cell Signalling, 12480S), anti-vimentin (1:1000, Proteintech, 10366-1-AP), anti-keratin14 (1:1000, BioLegend, 906004), anti-CD49f (1:1000, Sigma-Aldrich, ZRB1168) and anti-paxillin (1:1000, Invitrogen, PA5-34910); and visualised by probing with the following secondary antibodies (all at 1:10000 concentration): goat anti-guinea pig Alexa Fluor 594 (Jackson ImmunoResearch, 706-585-148), goat anti-rabbit Alexa Fluor 750 (Invitrogen, A-21039), goat anti-chicken STAR RED (Abberior, STRED-1005), and phalloidin (for actin staining, pre-conjugated to Alexa Fluor 555, Invitrogen, A34055). Cell nuclei were counterstained with Hoechst (1 µg/ml, Invitrogen, H1399).

For Western blot analysis, to allow for more surface area for the cells to grow to ultimately harvest a higher yield of protein, cells were plated in 6-well plates (Fisher Scientific, 140685). The same protocol and steps described above for the CenB-TGF*β* treatment were also performed for WB sample preparation, scaling the media and treatment used for the volume of a 6-well plate well. At day 0, day 7 and day 3 - as an added control of activation of EMT transcription factors - after addition of TGF*β,* cell monolayers were washed in ice-cold 1x PBS and 100µl of sample buffer - containing 8% w/v (g/100ml) SDS (Serva, 20765.01), 200 mM Tris-HCl pH 6.8 (SigmaAldrich, T1503), 40% v/v (ml/100ml) Glycerol (VWR International GmbH, 24384.290), 100 mM DTT (Biomol, 04010.10), Bromophenol Blue (Bio-Rad, 1610404) - was directly added to the plate. Cells were mechanically scraped and the lysate was collected into an eppendorf tube and boiled for 10min at 95°C. Lysates were then stored at -20°C until all samples were collected and used for further processing. WB was performed according to standard procedures. Briefly, gel electrophoresis was performed by running samples on a precast polyacrylamide NuPage 1.5mm 15 well 4-12% Bis-Tris gel (Invitrogen, NP0336PK2) with MOPS buffer (Invitrogen, NP0001), according to manufacturer’s instructions, in a XCell SureLock Mini-Cell system (Invitrogen, EI0001). Samples were run at constant 120V for 2h Gels were removed from their plastic cases, the stacking component excised, and re-equilibrated in 1X Novex Tris-Glycine Transfer Buffer (diluted from a 25X stock, Invitrogen, LC3675). In the meantime, a 0.45µm PVDF membrane (Millipore, IPVH00010) was activated in 100% Methanol for 15 seconds and placed in Milli-Q grade water for 2min; the membrane was then equilibrated in transfer buffer for at least 5min. Gel and membrane were arranged in a sandwich with the components of the XCell II Blot Module (Invitrogen, EI9051), according to manufacturer’s instructions. Wet WB transfer was performed at constant 95V for 2h. After transfer, the membrane was blocked in 5% milk in 1x 0.1% PBS-T (Frema) for 1 h at RT shaking. The membrane was then cut into 5 different strips at selected molecular weight, and stained shaking overnight at 4°C with the following concentration of primary antibodies diluted in 1% milk in 1x 0.1% PBS-T: 250 kDa Fibronectin (1:1000, Proteintech, 15613-1-AP), 130 kDa N-cadherin (1:1000, Proteintech, 22018-1-AP), 55 kDa **α**-tubulin (1:1000, Abcam, ab18251), 35 KDa GAPDH (1:1000, Cell Signalling, 2118), 29-35 kDa Snai1 (1:1000, Proteintech, 13099-1-AP), and 15 kDa Histone 3 (1:2000, Proteintech, 17168-1-AP). Membrane strips were washed three times in 1x 0.1% PBS-T for 10min each and probed with HRP-conjugated secondary antibodies against mouse or rabbit (1:5000, Jackson ImmunoResearch, goat anti-mouse: 115-035-068, goat anti-rabbit:111-035-046) for 1h at RT in the dark. Following incubation, membranes were washed again three times for 10min each with 1x 0.1% PBS-T and then developed using the Clarity Western ECL Substrate kit (BioRad, 1705060) according to manufacturers’ instructions and visualised on a Azure 280 imager (Azure BioSystems Inc.). Resulting images were processed in Fiji/imageJ.

#### IF/U-ExM image acquisition

For all IF experiments described above, except for the 5 color CenB-TGFβ experiment and paxillin experiments, Z-stack multiplex imaging was performed on the Zeiss LSM980 with AiryScan2, with a motorised stage with Piezo, via the ZenBlue software (v3.9, Carl Zeiss Microscopy GmbH) using a Plan-Apochromat 63x/NA 1.4 Oil DIC objective (Carl Zeiss Microscopy GmbH) and in multiplex SR-8Y mode, and laser lines: 405 nm (diode; 15 mW), 488nm (diode; 13 mW), 561nm (DPSS; 13 mW), 639nm (diode; 10 mW). 3D raw datasets were post-processed using the internal ZenBlue AiryScan deconvolution ‘AiryScan processing’ batch set to ‘Auto Filter’. Zeiss980 LSM confocal with AiryScan was also used for imaging U-ExM samples: a LD-LCI Plan-Apochromat 25x/0.8 multi-immersion autocorr. lens was used to localise the expanded sample and 140nm-spaced z-stacks of centrosomes were acquired with a C-Apochromat 40x/1.2 water autocorr. FCS M27 objective by scanning in AiryScan SR-mode. The same deconvolution algorithm as above was applied for U-ExM samples. For 5 color CenB-TGFβ IF experiments, z-stacks multiplex were acquired on a Leica Stellaris8 confocal microscope using a HC PL APO CS2 63x/NA 1.40 Oil and aided by a tunable 440-790 nm White Light Laser with adjustable spectral detection range 410nm-850nm and 4x HyD detectors. Post acquisition, cross talk correction (CTC) was performed using the LAS-X software (Leica Microsystems) to minimise the overlap between the signals of phalloidin conjugated to Alexa Fluor 555 and **α**/*β*-tubulin in Alexa Fluor 594. Dye separation was performed by estimating the distribution coefficient of the fluorophores, according to software guidelines, and all z-stack images of actin and **α**/*β*-tubulin were adjusted with the same calculated value matrix. Only CTC images were used for downstream analysis. Each set of experiments was acquired by maintaining the same microscope settings to allow for technical replicates. Z-stack imaging of CenB-TGFβ experiment of paxillin staining was performed on a Leica TCS SP8 DLS confocal using a 63x/NA 1.40 oil objective and laser lines 405nm, 488nm, 552nm, 638nm and 2x HyD detectors.

#### Image Analysis: ilastik, CellProfiler, CellposeSAM and more

Image analysis was performed using a mix of the open-source platforms Fiji/ImageJ v.1.54p (Schindelin *et al*, 2012), ilastik v1.4.1.post1 (Berg *et al*, 2019), CellProfiler (CP) v4.2.8 (Stirling *et al*, 2021) and Cellpose-SAM (Pachitariu *et al*, 2025), proprietary software Numbers v14.1 (Apple Inc., Cupertino) and GraphPad Prism v10.6.1 or above (GraphPad Software, Boston), and complemented by self-written Python code.

#### IF quantification of centriole numbers in MCF10A and MDA-MB-231

3D AiryScan processed images of MCF10A or MDA-MB-231 cells treated with CenB or TGFβ were analysed using Fiji/imageJ. Maximum intensity projections (MIP) were used to count the number of centrioles, identified as foci of CEP192 and colocalising with the *γ*-tubulin signal per nuclei, using the in-built Cell Counter plugin and cells were scored as either having 0, 1, 2 or >2 centrioles per nuclei, and results were plotted on a bar plot.

#### IF quantification of CenB treatment on MDA-MB-231

3D AiryScan images of CenB treated triple negative BC cells MDA-MB-231 were converted to MIP via Fiji/ImageJ. Cell segmentation was performed using cellpose-SAM: segmentation was performed on two channels (microtubules and nucleus) and by inputting the following parameters/values: flow_threshold = 0.6, cellprob_threshold = 0.8, tile_norm_blocksize = 100, diameter = 340, niter = 2000m batch_size = 32. Segmented cells touching the border were excluded and output masks were saved as TIFFs. MIP files of the multichannel fluorescence dataset alongside the cellpose-segmented cell masks were fed into the CP software. Each image was split into grayscale channels and the masks were imputed as objects in the Metadata tab. A custom pipeline was built using a combination of modules and parameters applied to the whole dataset. Nuclei segmentation was performed within the CP pipeline. This pipeline is publicly available under the name CenB_MDA-MB-231_D3_CP (see Data and Code Availability). Cells were manually scored for containing no centrosome or a single centrosome: cells with one centrosome/centriole present were excluded from the analysis. Features of cell body/nuclear shape and size (Calibrated Area, Solidity and FromFactor) collected from the cell body/nuclei objects were plotted as violin plots; while cell body features were used for PCA analysis.

#### U-ExM quantification of centriole length/volume in MCF10A

Length and volume of 3D-processed AiryScan images of centrioles in MCF10A cells, treated with *TGFβ* for 4 hours or 10 days, were analysed using ilastik. Single channel images of the acetylated tubulin signal were used in a pixel classification workflow followed by the object classification workflow. Approximately 10% of the complete dataset of centriole images was used to train the pixel/object classifications machine learning algorithm. For both pixel and object classification all possible training features were selected, and object classification was further refined with the following parameters/values during training: method=Simple, Smoothing: 1.00, Threshold: 0.35, Size Filter: 2500-100000 (min-max). Once the sparse annotation training was complete, and a user-defined satisfactory semantic segmentation generated, all images were processed in batch mode. Resulting probability maps were fed into the object classification workflow alongside their respective RAW image. Images for object classification were also processed in batch mode after training, resulting in object predictions and features table exports. Resulting lengths and volume measurements were spatially calibrated (multiplied by pixel width, pixel height and, for volume, voxel depth), and divided by the expansion factor mean of 4.00, to obtain µm measurements of biological relevance. Centriole length variation per conditions were plotted as violin plots.

#### IF quantification of TGFβ-induced EMT characteristics in MCF10A

3D AiryScan processed images of MCF10A cells, treated with TGF*β* for 4 hours or 10 days, were first converted to maximum intensity projections in Fiji/ImageJ. Cellpose-SAM segmentation was performed as described above for other AiryScan datasets. Cellpose-SAM segmented cells and RAW MIP images were processed in CP, following the same metadata/names&types steps as above and modifying the pipeline accordingly. Nuclei segmentation was performed with modules of the CP pipeline. The pipeline is publicly available under the name EMT-TGFb_MCF10A_T4-D10_CP (see Data and Code Availability). All measurements calculated for the nuclei object and the cell objects and collected from the modules were exported to a .csv file for subsequent plotting and analysis: cell body features were used for PCA analysis, while features of nuclear shape and size (Calibrated Area, Solidity and Extent) were plotted as violin plots.

#### IF quantification of Golgi apparatus features of Cen-TGFβ-treated MCF10A

3D AiryScan processed images of CenB-TGF*β-*treated MCF10A cells stained with the Golgi marker GM130 were converted to MIP using Fiji/ImageJ. Cell segmentation was performed as described above via cellpose-SAM, inputting the same parameters/values. CP analysis was also performed similarly to above, by variating the pipeline modules to identify the Golgi apparatus as one within each segmented cell. This pipeline is publicly available under the name CenB-TGFb_MCF10A_D0-D17_golgi_CP (see Data and Code Availability). Features of the cell body analysis (calibrated bounding box area and number of children) were plotted as violin plots.

#### IF quantification of EMT features of CenB-TGFβ treated MCF10A

Cross-talk corrected z-stack images of MCF10A cells, treated with the combinations of CenB and TGF*β* for 0 and 7 days, were first converted to MIP with Fiji/ImageJ. Cells were segmented by using cellpose-SAM as described above but by inputting nuclei, actin and microtubule channels and the same parameters/values. Nuclei segmentation was performed within the CP pipeline. Two separate pipelines were composed to analyse cells and nuclei. RAW images and corresponding cell segmentation masks were added to the CP tool with images as grayscale single channel images, and masks as objects, following the same metadata/names&types steps as above and modifying the pipelines accordingly. The pipelines are available under the names CenB-TGFb_MCF10A_D0-D17_CTC_CP_nuclei OR_cell (see Data and Code Availability). All parameters exported for cell body and nuclear objects were further analysed using Numbers and GraphPad Prism, and the UMAP workflow (see below). Features of cell body/nuclear shape and size (spatially calibrated area, solidity, and FromFactor) collected from the cell body/nuclei objects were plotted as violin plots. MeanIntensity features were calculated for the vimentin and keratin14 channels, as these are easily contained with each cell object, and plotted as scatter bars with depicted mean. For analysis of actin and microtubule, a first masking of the signal was performed with the use of ilastik. Pixel classification was performed on each channel independently, and the resulting probability masks were used to calculate microtubule directionality and actin/microtubule cross alignment based on structure tensor analysis. Results of our custom made analysis were compared to the output of AFT − Alignment by Fourier Transform workflow (Marcotti *et al*, 2021) and found to be equivalent in trend. The code for the cytoskeleton analysis is available under the name CytoActin-MT_analysis_FullPipeline.py (see Data and Code Availability).

The UMAP dimensionality reduction technique was generated by following the code and guidelines from the original paper’s depository (McInnes *et al*, 2018) (available at: https://github.com/lmcinnes/umap). The full feature table outputted from the CP cell object analysis was used as the input dataset, without any filtering. The main hyperparameters were set with the following values: n_neighbors=15, min_dist=0.15, n_components=2, and metric=’euclidean’. UMAP plotting was performed via the umap.plot package (v.0.4dev), also hosted at the above github depository.

#### IF quantification of paxillin/actin co-localisation

Four selected slices close to the coverslip of z-stack confocal images of CenB-TGFβ*-*treated MCF10A cells, stained for paxillin and actin, were converted to MIP with Fiji/ImageJ. MIP were split into single paxillin and actin channels and analysed using the Coloc2 plug-in in Fiji/ImageJ. We used the Manders’ and Pearson’s coefficients to establish, respectively, the co-occurance or fraction of overlap (‘the extent to which the signal from both molecules appears in the same pixels’) and the correlation (‘relationship between the variation in concentration of the two molecules’) between the paxillin and actin signals (Adler & Parmryd, 2021). Both the Pearson’s coefficient and the Manders’ thresholded M2 (tM2) coefficient were calculated to measure the correlation and co-occurrence (independent of signal intensity) between paxillin and actin, respectively, and plotted as box and whisker plots.

#### Cell division assay: live imaging and analysis

Total nuclei number change per frame over time was calculated as a proxy for cell division. For live imaging, MCF10A cells were plated in a 4-well glass-bottom chamber slide system (ThermoScientific, 177399) and subjected to the CenB-TGFβ protocol and treatments described above in detail. At day 0 and day 6, cells were washed and incubated in DMEM/F-12 without phenol red (Gibco, 21041025) - supplemented with only 2% horse serum and 100 ng/ml cholera toxin-containing the nuclear stain SiR-DNA-650 (1:1000, Spirochrome, SC007) and the microtubule stain SPY-Tubulin-555 (1:1000, Spirochrome, SC203). Dyes were allowed to intercalate into the cells for 2 hours before cells were washed, supplemented with fresh imaging media and mounted on the microscope stage for imaging. Live time-lapse imaging was performed on a Nikon-TiE widefield system run by NIS-Elements Advanced Research software (v4.60, Nikon Instruments Inc., Japan), fitted with a custom made incubator, equipped with a Hamamatsu Orca Flash 4 V2 camera and a LUMENCOR SPECTRAX lamp for illumination. Multichannel, multiposition images of all treatments were acquired every 15 min for 20h with a CFI P-Apo Lambda 20x/0.75 air objective. Time-lapse images were analysed via a combination of Fiji/imageJ, cellpose-SAM, ilastik, and python. The channel containing only the SiR-DNA-650 signal was used to segment the nuclei with cellpose-SAM, by setting the algorithm’s parameters/values to: flow_threshold = 0.6, cellprob_threshold = 0.8, tile_norm_blocksize = 100, diameter = 30, niter = 1000, batch_size = 32. Nuclei masks touching the border were discarded and the remaining masks were saved as new TIFF files. Raw fluorescent images of the timelapses alongside their respective nuclei masks were loaded into ilastik’s tracking pipeline, to obtain individual nuclei ID and tracking features. Total nuclei number per frame over time was extracted from the resulting exported .cvs file.

#### Plotting and Statistical Analysis

Except for the plots detailed below in this paragraph, all other graphs (bar, violin, PCA, etc.) were plotted using a combination of Numbers and GraphPad Prism, and statistical analysis was performed using the inbuilt functions in GraphPad Prism. Except where otherwise indicated in the figure legends, an ordinary two-way ANOVA followed by a Tukey’s post hoc multiple comparison test was applied for significance detection. MA plots, heatmaps and violin plots of the RNA-seq differential expression were plotted using RStudio and the Shiny app package. 3D PCA was plotted in python using the matplotlib library.

#### CenB-TGFβ RNA-seq: RNA isolation, library preparation and sequencing

Total RNA was extracted from MCF10A cells by using TRIzol Reagent (ThermoScientific, 15596026) according to the manufacturer’s instructions. Briefly, cells were seeded on 6-well plates and subjected to the CenB-TGFβ treatment as described in detail above. At the specific time points, cell monolayers were lysed and homogenized in 500 µl TRIzol, and the total RNA was collected from the aqueous phase after chloroform addition. RNA samples were then precipitated in isopropanol containing 10 µg glycogen for a better RNA pellet visualization, washed in 75% ethanol, resuspended in UltraPure RNase-free H_2_O (Invitrogen, 10977049), and incubated at 60°C for 10 min in a thermomixer. RNA quantity and purity/quality were determined on a high-throughput automated 4200 TapeStation System (Agilent, G2991BA), using the RNA ScreenTape Analysis kit (Agilent, 5067-5576), according to the manufacturer’s instructions. For library preparation, all samples were diluted to a final concentration of 300 ng/50 µl and placed, in triplicates, in a sealed and skirted 96-well plate. The libraries were prepared on a Beckman Coulter Automated Workstation Biomek i7 Hybrid (MC + Span-8) (Beckman Coulter, Inc., California, USA). For library preparation, an automated version of the NEBNext® UltraExpress™ RNA Library Prep Kit (New England Biolabs, E3330) was used, following section 1A - Express Protocol for use with NEBNext Poly(A) mRNA Magnetic Isolation Module (New England Biolabs, E7490). An adaptor dilution of 1 to 120 was used. The samples were individually barcoded using unique dual indices during the PCR using 16 PCR cycles. The individual libraries were quantified using the Qubit HS DNA assay (Thermo Fisher Scientific, Q32851) as per the manufacturer’s protocol. For the measurement 1 µl of sample in 199 µl of Qubit working solution was used. The quality and molarity of the libraries were assessed using Agilent Bioanalyzer with the DNA HS Assay kit (Thermo Fisher Scientific, Q32851) as per the manufacturer’s protocol. The assessed molarity was used to equimolarly combine the individual libraries into one pool for sequencing. The pool was loaded and sequenced on an Illumina NextSeq 2000 platform (Illumina, California, USA) using a P2 100 cycle kit, a read-length of 122 bp single-end reads and 650 pM final loading concentration. Library preparation, sequencing and analysis were performed by EMBL’s GeneCore facility.

#### RNA sequencing: read processing, alignment and quantification

Raw sequencing reads were processed using the nf-core/rnaseq pipeline version 3.21.0 (available at: https://github.com/nf-core/rnaseq/tree/3.21.0) (Ewels *et al*, 2020). Adapter sequences and low-quality bases were trimmed using Trim Galore v0.6.7. Reads were aligned to the human reference genome (GRCh38) using STAR v2.7.11b (Dobin *et al*, 2013) with the Ensembl genome annotation version 113. Gene-level quantification was performed using Salmon v1.10.3 (Patro *et al*, 2017) in alignment-based mode on the STAR alignments. Read counts and transcript length matrices were generated for downstream differential expression analysis.

#### RNA sequencing Differential Expression Analysis (DEA)

Differential gene expression analysis was conducted using the nf-core/differential abundance pipeline v1.5.0 (https://github.com/nf-core/differentialabundance/tree/1.5.0) with DESeq2 v1.44.0 (Love *et al*, 2014). Raw gene counts from Salmon were used as input. Genes were filtered to retain only those with a minimum abundance of 1 count in at least 1 sample. DESeq2 normalization was performed using the default size factor estimation with the ratio method. Dispersion estimation and statistical testing followed the Wald test with parametric fit type. Variance stabilizing transformation (VST) was applied for exploratory data analysis, including principal component analysis (PCA) and sample clustering using Ward’s hierarchical clustering method with Spearman correlation distances. Differentially expressed genes (DEGs) were identified using thresholds of |log2 fold change| ≥ 1 and adjusted *p*-value (Benjamini-Hochberg correction) < 0.05. For heatmap analysis and representation, genes from the Molecular Signatures Database (MsigDB v2025.1) Human Gene Sets (HALLMARK_EPITHELIA_MESENCHYMAL_TRANSITION v5.0, GO_EXTRACELLULAR_MATRIX v7.2 and HALLMARK_TGF_BETA_SIGNALLING v5.0) were used, and complemented with genes from selected publications (Taube *et al*, 2010; Vasaikar *et al*, 2021)

#### RNA sequencing: Gene Set Enrichment and Functional Analysis

Gene Set Enrichment Analysis (GSEA) was performed using the fgsea R package with 1,000 permutations, using gene-level statistics ranked by signal-to-noise ratio. The Hallmark gene sets from the Molecular Signatures Database (MSigDB v2025.1) were used for pathway analysis. Additionally, over-representation analysis was conducted using gprofiler2 v0.2.3 (Kolberg *et al*, 2020) with the human organism database (’hsapiens’), applying the g:SCS multiple testing correction method.

## Data and Code Availability

RNA-seq raw and processed datasets have been deposited in the EMBL-EBI BioStudies/European Nucleotide Archive (ENA) via Annotare under accession number E-MTAB-17456. Light microscopy image datasets, CellProfiler/cellpose/ilastik/python scripts and any other information are available upon request to the corresponding authors.

## Author Contributions

MLP, JMT, IH and NB conceptualised and designed the project. MLP, JMT, FJ, MB and NB performed the experiments and analysed the data. MLP and NB wrote the manuscript with contributions from all other authors.

## Acknowledgments

The authors acknowledge the support of the Advanced Light Imaging Facility (ALMF) for image acquisition, GeneCore for RNA sequencing, and the IT and HCP services for computational resources at EMBL Heidelberg. We would like to thank the Data Science Centre at EMBL, and in particular Charles Girardot and the Multimodal Open Data Integration Support (MODIS) team, for their valuable guidance in the management and submission of the RNA-Seq data. We would also like to thank the members of the Kreshuk laboratory, and in particular Dominik Kutra, for assistance in resolving ilastik-related issues and code supervision. This publication was supported by the European Molecular Biology Laboratory and through state funds approved by the State Parliament of Baden-Württemberg for the Innovation Campus Health + Life Science Alliance Heidelberg Mannheim.

## Extended View Data

**Figure. EV1.**
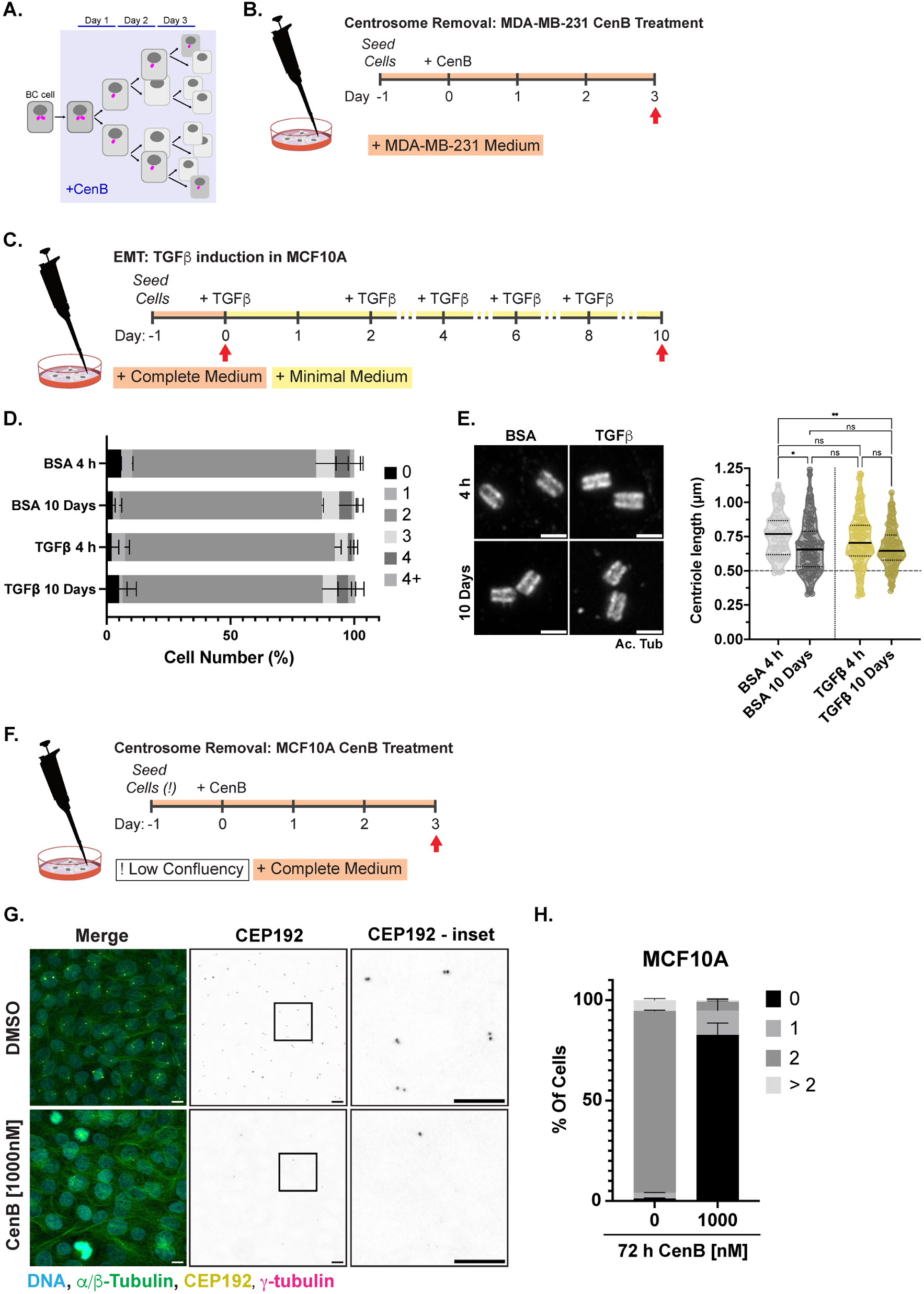
Centrinone B (CenB) depletes centrosomes in breast cells while EMT does not affect centriole ultrastructure nor numbers. **A.** Schematic of CenB mode of action on dividing cells: after each cell cycle (or days), an increasing percentage of daughter cells is left without a centrosome (magenta cylinders). **B.** Schematic representation of CenB treatment protocol to remove centrosomes in MDA-MB-231. Red arrows represent day/time of fixation. **C.** Schematic representation of TGFβ-stimulated EMT induction protocol in MCF10A cells. Red arrows represent day/time of fixation. **D.** Bar plot shows quantification of centriole number in MCF10A cells after TGFβ treatment for 4 hours (4 h) or 10 days. Bars show mean ± SD of two independent experiments with a minimum of N=70 cells per condition. **E.** Representative AiryScan confocal UExM images of centrioles from MCF10A cells treated with BSA (control) or TGFβ at 4 h and 10 days. Maximum intensity z-projections show acetylated tubulin signals. Scale bar represents biological value: 500nm (and equivalent expanded physical value: 2µm). Violin plot showing median and quartiles of the length variation of centrioles, measured with the software ilastik. Each data point represents one centriole, derived from two independent experiments with a minimum N=54. For statistical significance, in all violin plots, an ordinary two-way ANOVA followed by Tukey’s multiple comparison test was performed. *p-values* meaning: non significant (ns)=> 0.05; *=0.05; **=≤ 0.01. **F.** Schematic representation of CenB treatment protocol in MCF10A to remove centrosomes. Red arrows represent day/time of fixation. **G.** Representative AiryScan confocal IF images of methanol fixed MCF10A treated with DMSO or 1000nM CenB for 72 hours. Maximum intensity z-projections show merge of *α*/β-tubulin (green), CEP192 (yellow), γ-tubulin (magenta) and nuclei (DNA, Hoechst, cyan), and individual channel for CEP192 (grayscale, inverted). Inset represents zoom-in of selected region (boxed) of CEP192 (grayscale, inverted) channel to better visualise centrioles. Scale bars: 10µm. **H.** Bar plot shows quantification of centriole number in MCF10A cells after 72 h of DMSO or 1000nM CenB treatment. Bars show mean ± SD of two independent experiments with a minimum of N= 347 cells per condition.

**Figure. EV2.**
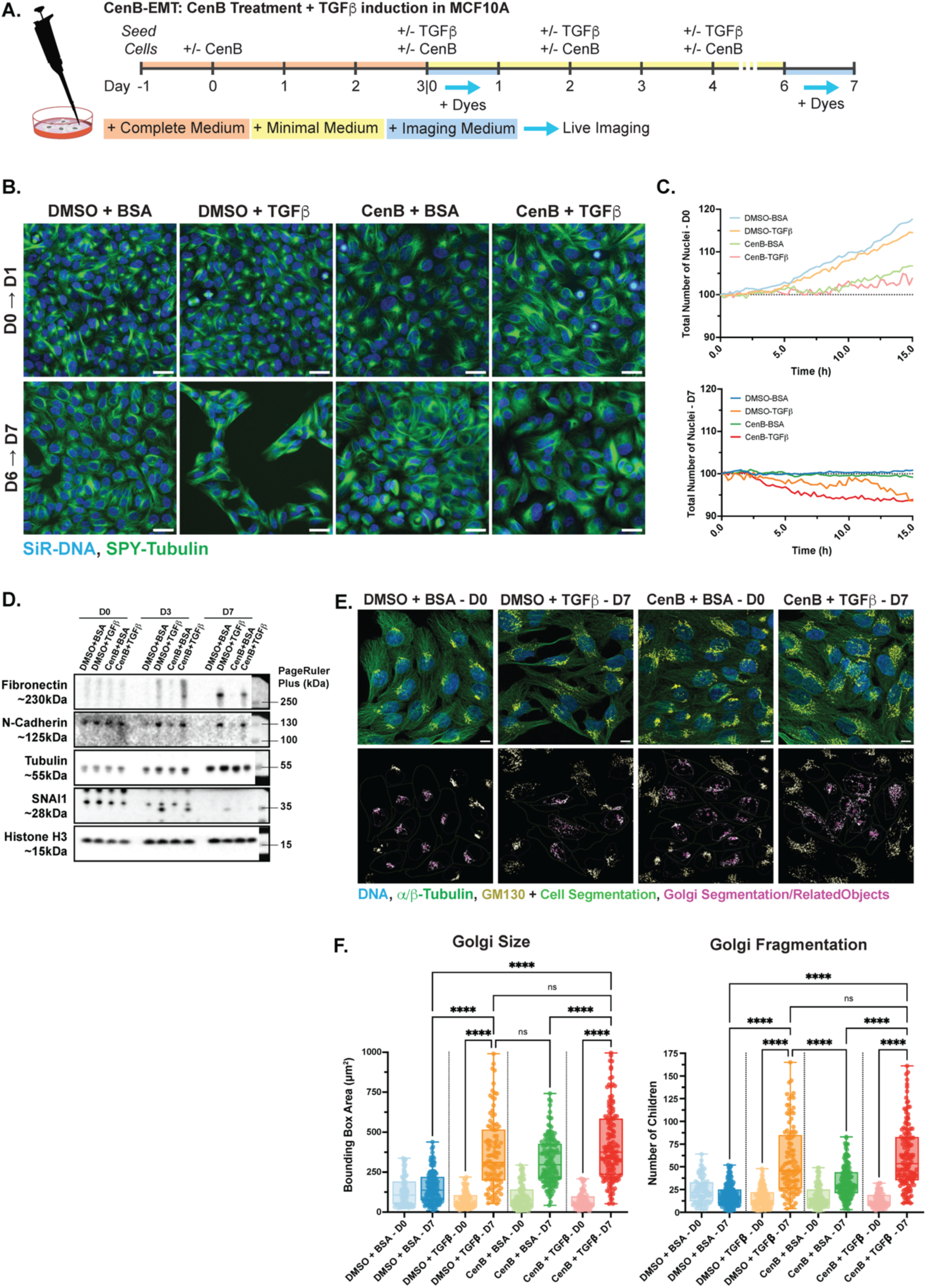
CenB impacts multiple organelles while cell cycle differences do not fully account for EMT delay. **A.** Schematic representation of CenB-treated cells followed by TGFβ induction protocol. Light blue arrows represent the overnight time period between days D0-D1 and D6-D7 where cells were subjected to time-lapse live imaging, after being incubated for 2 hours with live-dyes. **B.** Representative widefield frames of live time-lapse imaging of MCF10A cells treated with CenB-TGFβ at D0 and D7, in all combinations. Live dyes added SiR-DNA-650 and SPY-tubulin-555 highlight the nuclei and microtubules respectively. Scale bar: 50µm. **C.** Color coded quantification of percentage change of total nuclei number over time, as a proxy for cell division, for MCF10A cells treated with CenB-TGFβ in all combinations, at D0 (top graph) and D7 (bottom graph). Graph represents mean value per time-point and is derived from three independent experiments with a minimum of three acquired imaging locations per timepoint. **D.** Western blot analysis of CenB-TGFβ treated cells, in all conditions, at D0, D7 and day 3 (D3). Cells were further harvested at D3 to visualise presence of the transcription factor SNAI1, which is not active either at D0 or D7. Immunoblotting was performed against the EMT markers: fibronectin, N-cadherin, SNAI1, and **α**-tubulin and Histone H3 as loading controls. Right lane image: prestained PageRuler Plus molecular ladder ranging from 15-250kDa. Contrast was modified for visualisation purposes. Represented here one of two independent repeats. **E.** Representative AiryScan confocal IF images of methanol-fixed MCF10A cells with selected treatments: DMSO-BSA at D0, DMSO-TGFβ at D7, CenB-BSA at D0 and CenB-TGFβ at D7. Top panel: Maximum intensity z-projections of selected conditions show both merge of GM-130 (yellow), *α*/β-tubulin (green), and nuclei (DNA, Hoechst, cyan). Scale bar: 10µm. Bottom panel: maximum intensity projections of GM-130 channel also depicting the segmented cell bodies (green outline), the overall segmentation of the Golgi (yellow outline) and the bounding box and object identification number of the Golgi apparatus within each cell soma (magenta outline). **F.** Box and whiskers plot, showing min to max and median values, of bounding box area (in µm^2^) measurements of segmented Golgi (left graph) and number of children (right graph), calculated as masked objects within the bounding box area or grouped segmented Golgi. Measurements produced with CP analysis and derived from three independent experiments with a minimal N=83 Golgi per condition. For statistical significance in the box and whiskers plot, an ordinary two-way ANOVA followed by Tukey’s multiple comparison test was performed. *p-values* meaning: non significant (ns)=> 0.05; ****=≤ 0.0001. Only selected significance are shown, for a full description of significance see Expanded View Table Fig. EV2.

**Figure. EV3.**
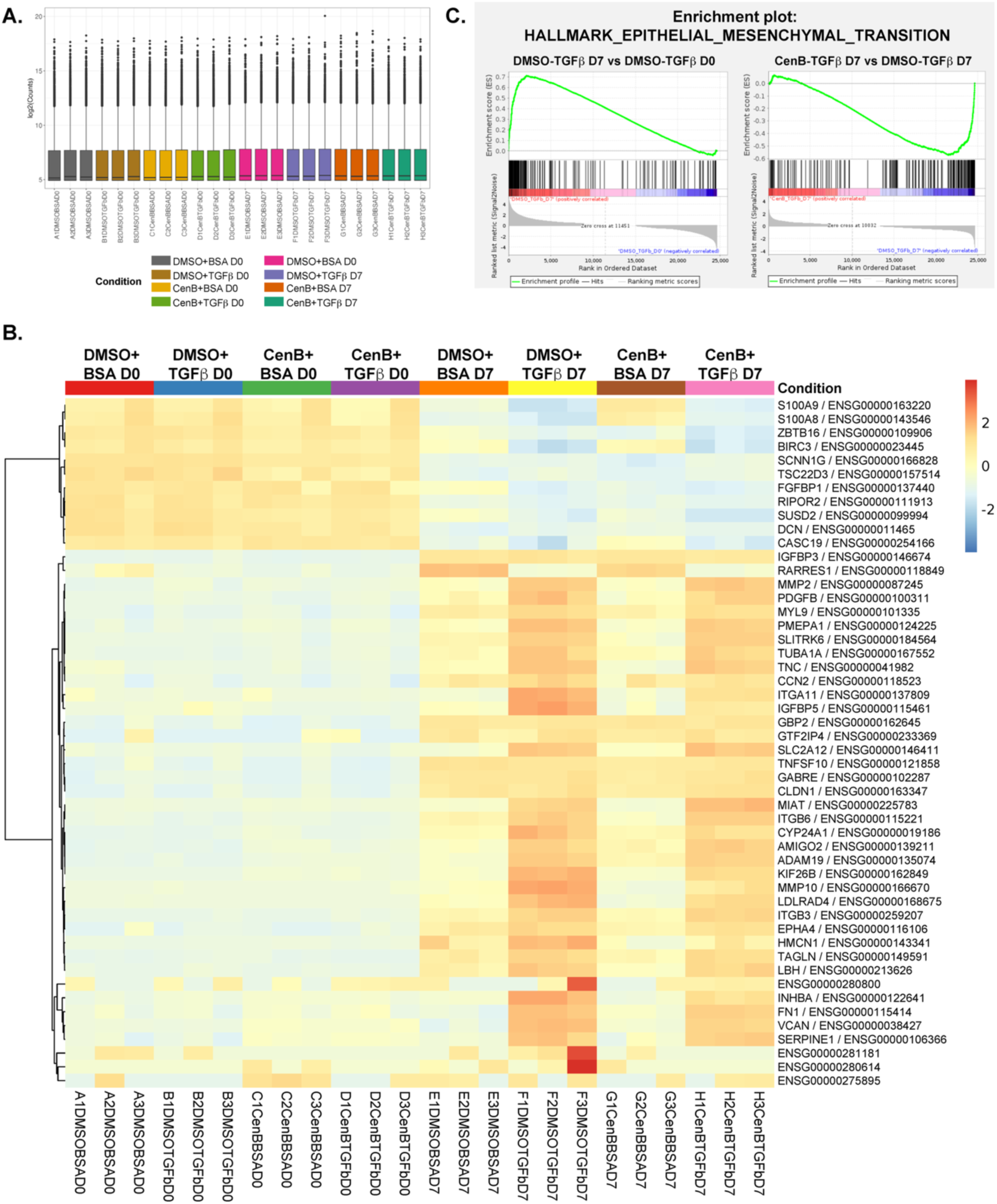
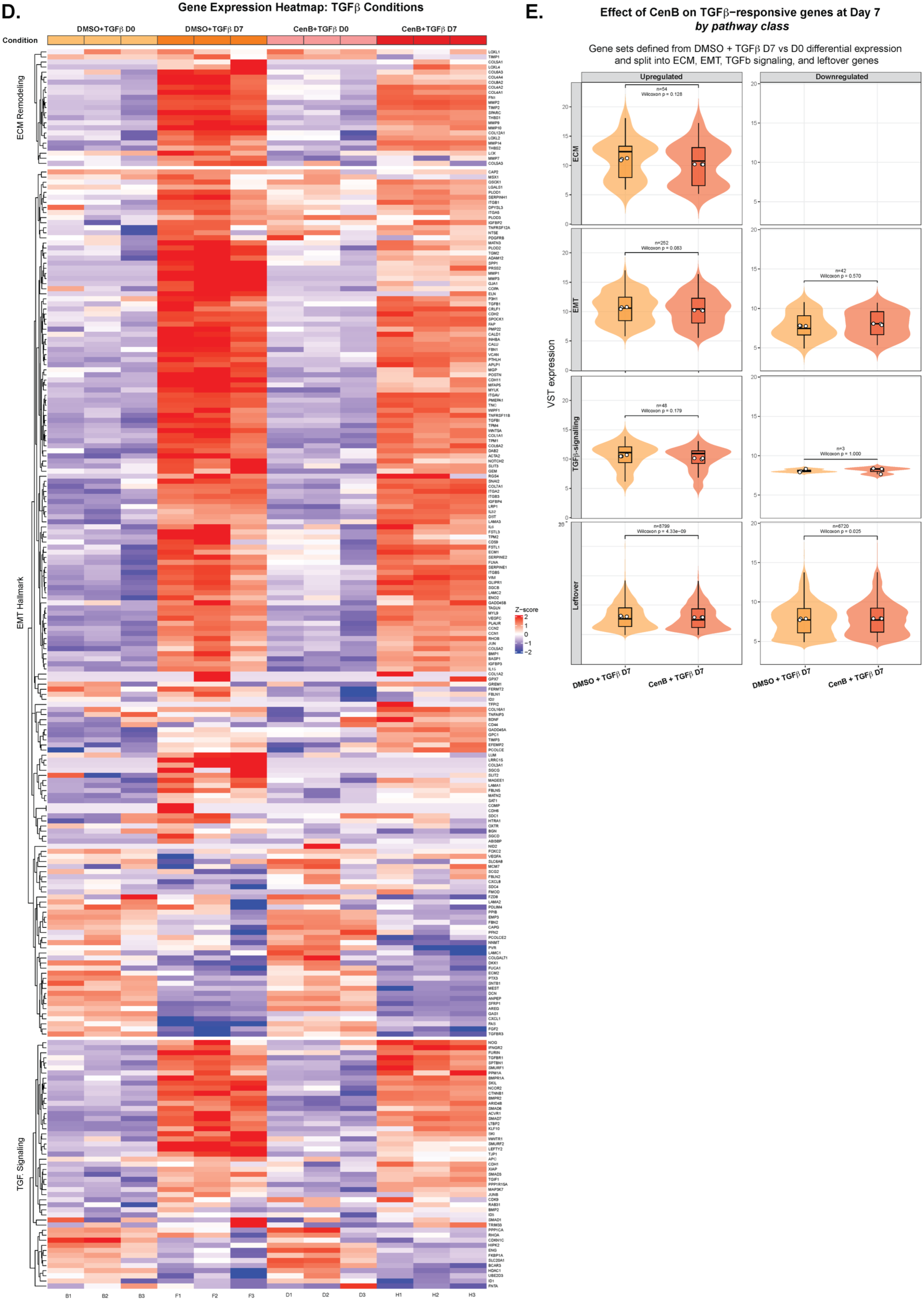
RNA-seq analysis supplementary datasets. **A.** Bar graph of correlation of quality of triplicate RNA samples for CenB-TGFβ treated MCF10A cells at D0 and D7 in all conditions. **B.** Heatmap of 50 most differentially regulated genes between MCF10A cells treated with CenB-TGFβ at D0 and D7 in all combinations. **C.** Gene Set Enrichment Analysis (GSEA) plot of EMT hallmark gene set, featuring opposite enrichment scores (ES) and leading edge-subsets for DMSO-TGFβ D7 vs D0 (left graph) and CenB-TGFβ vs DMSO-TGFβ at D7 (right graph). **D.** Heatmap of differentially expressed genes involved in the pathways of ECM remodelling, EMT hallmark and TGFβ-signalling for conditions treated with TGFβ before and after stimulation. Gene list selection was compiled from GSEA EMT hallmark as well as from selected references (See Material and Methods: *RNA sequencing Differential Expression Analysis (DEA))*. **E.** Violin plot representing the effects of CenB on TGFβ-responsive genes after TGFβ treatment by pathway class. Pathway class analysed from top to bottom: extracellular matrix (ECM), epithelial-to-mesenchymal transition (EMT), TGFβ-signalling and all remaining (Leftover) genes. Variance Stabilising Transformation (VST) expression levels are illustrated for upregulated (left side panels) vs downregulated (right side panels) genes, with the number of genes analysed per condition written above each significance score. For significance a Wilcoxon rank-sum test was performed and the *p-value* obtained specified.

